# Evaluating the Effects of Propiconazole on Hard Fescue (*Festuca brevipila*) via RNA sequencing and Liquid Chromatography–Mass Spectrometry

**DOI:** 10.1101/2020.11.17.387290

**Authors:** Yinjie Qiu, Dominic Petrella, Florence Sessoms, Ya Yang, Mark Esler, Cory D. Hirsch, Garett Heineck, Adrian Hegeman, Eric Watkins

## Abstract

Propiconazole is often used to remove fungal endophytes from turfgrass to study the effects of *Epichloë* endophytes. However, besides a fungicidal effect, propiconazole can bind to the genes in the cytochrome P450 family and affect the biosynthesis of brassinosteroids. For this reason, outside of fungicidal application, propiconazole has also been used as plant growth regulator. In this study, we used a combination of RNA sequencing and liquid chromatography–mass spectrometry (LC-MS) to study how hard fescue (*Festuca brevipila*) responded to the high dose of propiconazole treatment. To test the long-term effect of the heavy use of propiconazole on plants, we inoculated with *Microdochium nivale* (causal agent of pink snow mold) half year post the last fungicide application. Propiconazole-treated plants showed enhanced pink snow mold resistance. This study suggested that the high dose use of propiconazole fungicide resulted in phenotypic and physiological changes in the plant such as slow growth and change in disease resistance. Genes and pathways affected by propiconazole identified in this study provide turfgrass breeders new information for genetic improvement of hard fescue and also provide turfgrass management new ways to control turfgrass diseases.

## Introduction

Turfgrass diseases are one of the greatest challenges to turfgrass managers. Reduction of turfgrass diseases can be achieved through management methods such as increased air flow, reducing soil compaction, and the use of fungicides (Vargas 2018). Of these, fungicide applications appear to be the most effective way for common turfgrass disease control. For example, most golf course superintendents apply fungicides every 14 to 21 days during summer months to control dollar spot *Clarireedia jacksonii* on creeping bentgrass putting greens (Vincelli 2004). Vincelli et al. (2003) observed the best dollar spot management control when spraying triadimefon (0.38 kg a.i./ha) and chlorothalonil (8.0 kg a.i./ha) in 407 liter of water. For pink snow mold (*Microdochium nivale*) control, the use of five fungicides treatment showed the reduction of disease severity (Jung et al., 2007). The heavy use of fungicide application could control diseases, but creates issues such as fungicide runoff (Watschke, Mumma et al. 2000) and fungicide resistance (Jo, Won Chang et al. 2008). For these reasons, turfgrass breeding and genetics research has focused on breeding for disease resistance (Bonos and Huff 2013).

Turfgrass breeders can develop more resistant germplasm through either traditional breeding or transgenic approaches (Belanger, Bonos et al. 2004, Bonos, Clarke et al. 2006, Dong, Tredway et al. 2007). Additionally, bioagents such as fungal endophytes have also been exploited as a strategy to enhance biotic stress tolerance of turfgrass (Bacon, Richardson et al. 1997). For example, endophytes have been found to improve the biotic stress of turfgrass such as resistance to sod webworms (*Crambus* spp.) in perennial ryegrass (*Lolium perenne*) (Funk, Halisky et al. 1983), and dollar spot and causal agent resistance in fine fescues (*Festuca* spp.) (Bonos, Wilson et al. 2005, Clarke, White Jr et al. 2006). Fungal endophytes were also found to improve abiotic stress tolerance such as heat (Tian, Huang et al. 2015) and drought (White, Engelke et al. 1992). When studying how endophytes improve plants stress tolerance, most studies carry out a systemic fungicide application with high rates and multiple applications to eliminate endophytes in the turfgrasses to create an endophyte-free population (Hesse, Schöberlein et al. 2003, Kane 2011, Heineck, Watkins et al. 2018). Of the commonly used fungicides, propiconazole (cis–trans-1-[2-(2,4-dichlorophenyl)-4-propyl-1,3-dioxolan-2-ylmethyl]-1H-1,2,4-triazole) shows the best results for fungal endophyte removal (Harvey, Fletcher et al. 1982, Latch and Christensen 1982). Propiconazole is a systematic triazole fungicide that targets fungal 14-alpha demethylase enzyme (14-alpha demethylase inhibitor) from demethylating a precursor to ergosterol (Kwok and Loeffler 1993). Besides fungicidal effects, propiconazole targets the brassinosteroid (BR) biosynthesis pathway in plants through binding to CYP90B1, CYP90C1, and CYP90D1 to inhibit the side chain hydroxylation between campesterol and teasterone (Hallahan, Heasman et al. 1988, Asami, Mizutani et al. 2001, Ohnishi, Szatmari et al. 2006, Hartwig, Corvalan et al. 2012). The brassinosteroids are a group of important phytohormones that function in multiple aspects of plants such as growth and development, along with stress tolerance (Mandava 1988, Nakashita, Yasuda et al. 2003, Bajguz and Hayat 2009). Plants with deficient BR usually show a dwarf phenotype (Noguchi, Fujioka et al. 1999). The BR and gibberellin acid pathway work together to regulate plant growth and development (Li, Wang et al. 2012). For these reasons, the high rate application of propiconazole could confound any effects observed from the loss of fungal endophyte. Therefore, it is important to study both short- and long-term effects of high propiconazole application rates for the endophyte removal process.

In this study, we carried out RNA sequencing and untargeted metabolomics approach to study hard fescue responses to the propiconazole fungicide. In order to evaluate how high rates of propiconazole affected plant responses in the long term, pink snow mold was used to inoculate the plant to evaluate the fungicide effect.

## Material and Methods

### Plant Material and Sampling

A single genotype of *F. brevipila* (SPHD15-3), from the University of Minnesota turfgrass breeding program, which was used to generate a *F. brevipila* reference transcriptome (Qiu et al., 2020) was vegetatively propagated into individual 5-tiller plants in soilless media (BRK Promix soil, Premier Tech, USA). This genotype was previously confirmed as endophyte free using a commercial Phytoscreen Immunoblot Kit [ENDO797-3, Agrinostics, http://www.agrinostics.com]. Because a previous study showed the effect of propiconazole on creeping bentgrass (*Agrostis stolonifera* L.), we used a single genotype ‘Penncross’ creeping bentgrass as a positive control for the fungicide application and used the same propagation method described above to generate the plant population. All plant material was grown in the Minnesota Agricultural Experiment Station Plant Growth Facilities in St. Paul, MN under a 16-hour photoperiod with every-other-day watering and weekly fertilization using a liquid fertilizer solution (200 ppm N, 22 ppm P, 83 ppm K, 114 ppm S, 2.5 ppm Fe, 750 ppb Mg, 100 PPb B, 50 ppb Cu, 280 ppb Mn, 500 ppb Mo, and 81 ppb Zn).

### Fungicide Application

Fungicide application was carried out on plants with approximately 8 tillers using Kestrel Mex fungicide which has propiconazole as the active ingredient (Phoenix Environmental Care, USA). The fungicide was sprayed at the rate of 1.31 ml m^-2^ between 9 and 10 am at one-week intervals (a total of five applications) in the greenhouse starting at the end of February 2019. A total of five fungicide applications were made.

### Tissue Sampling

One week after the second fungicide application, leaf samples were harvested for transcriptome sequencing and metabolomics profiling. Briefly, 1-2 cm of leaf tissue that was 2 cm from the leaf tip was harvested and flash frozen into 1.5 mL microcentrifuge tubes for RNA extraction. The plant tissues that were 1-2 cm below the cutting site for RNA sequencing were used to for LC-MS (**Figure S1**). All samples were stored at −80°C prior to analysis. Prior to tissue sampling, data was collected for turfgrass quality (using a 1-9 scale with 9 = best turfgrass quality as described by Kran et al., (Krans and Morris 2007) and phenotypic changes including plant leaf color, plant size change, were recorded using a Nikon D500 camera.

### RNA Extraction and Sequencing

RNA extraction was done using Quick-RNA Miniprep Kit (ZYMO Research, Catalog number R1055) following the manufacturer’s instructions. The RNA was diluted in DNase/RNase-Free water and stored at −80°C. Samples with RNA Integrity Number (RIN) >7.5 were used for sequencing library construction. Illumina sequencing libraries were constructed using TruSeq^®^ Stranded mRNA Library Prep kit (Illumina) and sequenced on Illumina HiSeq 4000 instrument with 150 bp paired-end sequencing mode. All sequencing data generated from this study was deposited at NCBI under bioProject PRJNA606332.

### Illumina Sequencing Data Processing and *de novo* Assembly

To process Illumina raw reads, adaptor sequences were trimmed using the Trimmomatic program (v 0.32) with the default setting (Bolger, Lohse et al. 2014). Quality trimming was performed using seqtk tool with -q 0.05 (https://github.com/lh3/seqtk). Trinity (v 2.4.0) was used to assemble transcriptomes using sequencing reads generated from both fungicide and snow mold inoculation experiments (Grabherr, Haas et al. 2011). Trinity program parameters were set as max memory 200G, 24 CPU cores, and bflyCalculateCPU, with minimum contig size of 200 bp. To improve assembly quality, a previously published *F. brevipila* Pacbio Isoform sequencing transcriptome (Qiu, Yang et al. 2020) was used and combined with the *de novo* assembly generated in this study. To merge and remove redundant transcripts in the combined transcriptome, ‘cd-hit-est’ from the CD-HIT (v 4.8.1) package was used to cluster redundant sequences with 95% sequence identity (Li and Godzik 2006). To evaluate the completeness of the *F. brevipila de novo* and hybrid transcriptomes, we used 1335 core embryophyte genes (embryophyta_odb10) from Benchmarking Universal Single-Copy Orthologs (BUSCO v3) (Simão, Waterhouse et al. 2015).

### Transcriptome Functional Annotation

The combined transcriptome was blast searched against NCBI non-redundant (NR) protein and SwisProt, UniProt protein database using diamond blastx (v 0.9.13) with e-value < 1e-5. Kyoto Encyclopedia of Genes and Genomes (KEGG) pathway analyses were performed using the KEGG Automatic Annotation Server (KASS, https://www.genome.jp/kegg/kaas/). *Oryza sativa, Brassica napus, Zea mays, Arabidopsis thaliana*, and *Aegilops tauschii* were used as references with a single-directional best hit model. The TransDecoder pogram was used to generate transcriptome protein sequences to perform Gene Ontology (GO) search using interproscan program (Quevillon, Silventoinen et al. 2005).

### Gene Expression Analysis

To investigate the gene expression change between propiconazole- and water-treated plants, expected number of fragments per kilobase of transcript sequence per millions mapped reads (FPKM) was calculated using RSEM program (Trapnell, Williams et al. 2010, Li and Dewey 2011) and further processed using edgeR programs implemented in the Trinity pipeline (Robinson, McCarthy et al. 2010, Trapnell, Williams et al. 2010). Differentially expressed genes (DEGs) were determined using DE_analysis.pl script in Trinity pipeline, further filtered with a False Discovery Rate (FDR) of 0.05, and minimum log fold change of 2.0.

### Differentially Expressed Cytochrome P450 genes

Because of high sequence feature similarity within the P450 gene family, a two-step approach was taken to annotate CYP450 genes identified in this study. The first step used a phylogenetics method. Briefly, *A. thaliana* cytochrome P450 protein sequences were downloaded from “The Cytochrome P450 homepage” reported by D. R. Nelson (http://drnelson.uthsc.edu/CytochromeP450.html) (Nelson, Schuler et al. 2004). Pseudogene sequences were removed and the remaining named *Arabidopsis* P450 sequences were used as a reference. Because the CYP92 gene family was lost in *Arabidopsis* lineage (Nelson, Schuler et al. 2004), protein sequences of eight genes within CYP92 family were downloaded from the Rice Genome Annotation Project (http://rice.plantbiology.msu.edu) (**Table S1**). CYP protein sequences of the differentially expressed CYP450 genes were added to the P450 references prior to performing phylogenetic relationship reconstruction. Protein sequences were aligned using MAFFT with-auto option (Katoh and Standley 2013). The alignment was inspected using Jalview program (v 1.0). Sequences before the starting codon were trimmed using TrimAl (Capella-Gutiérrez, Silla-Martínez et al. 2009). To construct the maximum likelihood (ML) tree, the best-fit model was predicted using iqtree (v 1.6.11) with -st AA -m TESTONLY option (Kalyaanamoorthy, Minh et al. 2017). The model with the best Bayesian Information Criterion (BIC) was selected to reconstruct phylogenetic tree via iqtree (v 1.6.11) with 1000 bootstrap replications (Nguyen, Schmidt et al. 2014). The phylogenetic tree was visualized using FigTree (v 1.4.3) (https://github.com/rambaut/figtree) and color coded by CYP Clan (Rambaut 2012). The second step was done by blasting protein sequences of candidate genes via NCBI blastp.

Top hit (Max score) annotation that was consistent with the phylogenetic results was considered as the final annotation for the transcripts.

### Untargeted Metabolomics Analysis

Propiconazole-treated plants were also analyzed for metabolomic changes compared to water-treated plants. To improve the accuracy of the LC-MS run, 4-5 subsamples were analyzed for each biological replicate for water and propiconazole treatments. Samples were extracted in 90% aqueous methanol (Sigma-Aldrich HPLC Plus grade methanol and Fisher Scientific Optima™ water) using 10 *μ*L solvent per mg of fresh-frozen tissue. After suspending samples in solvent, microcentrifuge tubes were placed back into a −80° C freezer for 48 hr. One 5-mm tungsten bead was added to each sample tube, and samples were ground in a bead mill (Genoginder SPEX SamplePrep) for 12 mins (1,400 RPM for 2 mins, and 700 RPM for 10 mins) in −20°C grinding blocks. Samples were vortexed and centrifuged at 14,000 g for 10 mins (4°C), and the supernatant was transferred into new tubes and re-centrifuged. To remove chlorophyll from the supernatant, all samples were diluted 1:2 in Optima™ water and were placed at −80° C overnight. Diluted extracts were then centrifuged at 16,000 g (4°C) for 30 mins. Final extracts were aliquoted into glass autosampler vial inserts, which were then inserted into autosampler vials for LC-MS analysis. A quality control pooled sample was also made using aliquots of extracts from all samples.

Untargeted metabolite analysis was performed using both reverse phase and normal phase methods. Both pool and blank samples were sampled four times throughout the run as quality controls. The reverse phase was performed using Waters Acquity UPLC HSS T3 1.8 Microm, 2.1 x 100 column (C18), and normal phase was performed with a Millipore SeQuant ZIC-cHILIC 3um, 100A, 100 x 2.1 mm column (cHILIC). Analyses were performed using an UltiMate 3000 High Performance Liquid Chromatography (ThermoFisher), and Q Exactive Hybrid Quadrupole-Orbitrap Mass Spectrometer (ThermoFisher).

For the C18 column run, the Thermo UltiMate 3000 UPLC pump, auto sampler, column compartment, and diode array detector (DAD) were set with the following parameters: flow rate = 0.4 mL per min; 10% B (0.1% Formic acid in acetonitrile) 1-2 min, 10-95% B; 2-25 min 95% B; 25-28 min 95-10% B 28-30 min; 210, 254, 340, and 520 nm were analyzed using the DAD along with full scan UV-vis; column temperature=40°C. The mass spectrometer parameters were set as: probe height = C; Full MS Scan range = 200 - 1500 m/z; Positive and negative ionization were monitored simultaneously using polarity switching; Sheath gas flow rate = 50; Aux gas flow rate = 20; Sweep gas flow rate = 1; Spray voltage = 3.8 kV; Capillary temp = 350 °C; S-lens RF level = 50; Aux gas heater temp = 300 °C.

For the cHILIC run, the Thermo UltiMate 3000 UPLC pump, auto sampler, column compartment, and diode array detector (DAD) were set with the following parameters: flow rate = 0.4 mL per min; 98% B hold 0-2 min; 98-55% B 2-32 min; 55-95% B 32-34 min; 95% B hold 34-37 min; and the UV-vis = NA; column temperature=40°C. The mass spectrometer parameters were set as: probe height = C; Full MS Scan range = 50 - 750 m/z; positive and negative ionization were monitored simultaneously using polarity switching; sheath gas flow rate = 5 Aux; gas flow rate = 0; sweep gas flow rate = 0; spray voltage = 3.3 kV; capillary temp = 320 °C; S-lens RF level = 55 Aux; gas heater temp = 0.

### Liquid Chromatography-Mass Spectrometry Data Processing

LC-MS data processing was done using the Work4Metabolomics based on (galaxy.workflow4metabolomics.org, version 3.3) for untargeted data analysis. Based on polarity, raw data files were split into positive and negative mzML files using the msconvert program (ProteoWizard release: 3.0.11252). The workflow for the data processing was done following instructions by Galaxy training for mass spectrometry: LC-MS analysis (https://galaxyproject.github.io/) with adjusted bandwidth for peak identification parameter. Detail preprocessing and postprocessing workflow is shown in **Figure S2**, and **Figure S3**. LC-MS features were used for unsupervised multivariant analysis and scatter plots were produced with PC1 and PC2 data to evaluate differences in water and propiconazole treatments. The identity of the top 10 metabolite features that were driving principal component (PC) 1 for positive and negative phase of both columns were predicted by searching accurate mass m/z data via KEGG compound and the human Metabolome Database (HMDB, hmdb.ca/spectra/ms/search), and Metlin.

### Long-Term Effect of Propiconazole on *F. brevipila*

To study the long-term effects of high rates of propiconazole on *F. brevipila*, we inoculated a pink snow mold isolate PSM to both propiconazole-treated and untreated plants to evaluate the disease severity (isolate was obtained from Dr. Paul Koch, University of Wisconsin, Madison). To avoid the effect from residual fungicide propiconazole, we carried out two rounds of vegetative propagation. Briefly, one month after the fifth fungicide treatment, single tillers were split from the original plant and were transplanted into fresh soilless media. The newly transplanted tillers were grown in the greenhouse in the condition described above for four months. Tillers of these transplants were split and transplanted in fresh soilless media again and were grown for one month to reach approximately five tillers before performing snow mold inoculation. Therefore, the analyzed plant had experienced the final propiconazole applications six months prior to inoculation. The scheme of the plant material preparation was shown in **Figure S4**.

### Inoculum Preparation

*Microdochium nivale* inoculum was prepared following the protocol described by Tronsmo (1993), Hofgaard (2006), and Kovi (2016) with modification. Briefly, the isolate was first cultured on potato dextrose agar (PDA) and incubated at 12 °C in the dark. Agar plugs (5 mm in diameter) were exercised from the PDA plate, transferred in a flask that contained 50 mL potato dextrose broth (PDB) and were incubated at 12 °C. After 15 days of growth, the mycelium was harvested by filtering through cheesecloth and homogenized in distilled water containing 0.01% TWEEN 20 (Sigma) using PRO Scientific Bio-Gen PRO200 Homogenizer. The inoculum was diluted to an optical destiny of 0.5 at 430 nm prior inoculation.

### Pink Snow Mold Inoculation

Plants were transferred into a dark walk-in cooler set at 10 °C one hour prior to inoculation. Inoculation was performed by dropping 1 mL inoculum to the crown region for the treatment group, and for mock-treated plants water drops were used as the control. Moist paper towels were used to cover plants to mimic snow cover and maintain moisture. One day post inoculation, plant tissues (1-3 cm above ground, **Figure S5**) were harvested for RNA sequencing for gene expression analysis (three biological repeats each of propiconazole-pretreatment plant with/without snow mold and water-pretreatment plant with/without snow mold). Sequencing and DEG identification were carried out using the method described earlier. Fourteen days post inoculation, the maximum mycelium spread length on the leaf (between crown region) was measured with a ruler to compare the different disease severity.

## Results

### Plant Phenotype Observation after Propiconazole Treatment

One week following the fifth propiconazole application, the positive control creeping bentgrass plants showed wider and thicker leaf tissue along with stunted plant growth as expected (**Figure 1 A, B**). *Festuca brevipila* had a similar response with slower growth and darker leaf color with propiconazole treatment (**Figure 1 C, D**) (photos were taken from the same height). One month after the last fungicide application, propiconazole-treated *F. brevipila* had smaller plant size compared with the control groups (size rating score of 7.67±0.58 for treated plants vs 9.0 ± 0.00 for non-treated).

**Figure 1.**
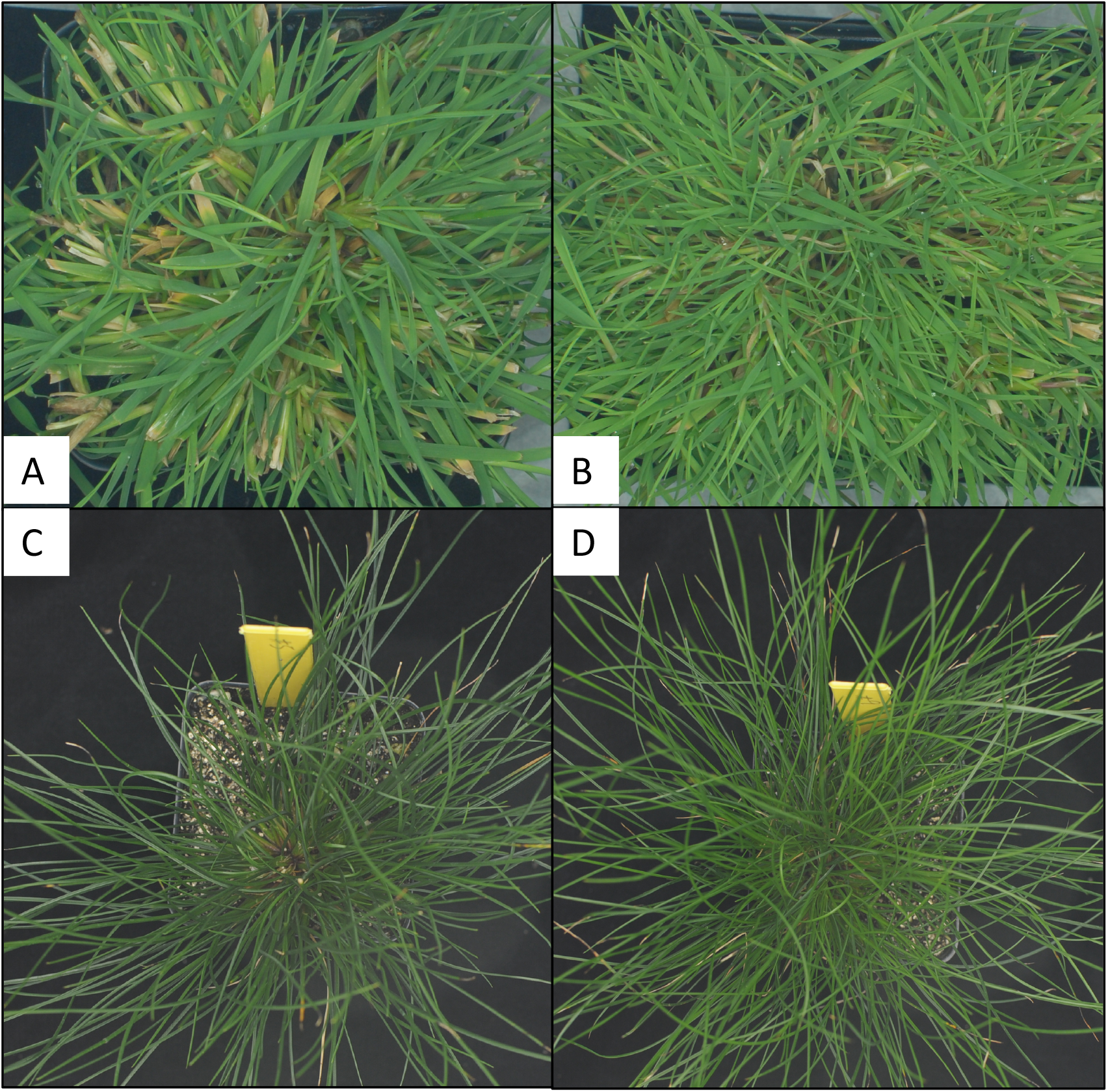
*Agrostis stolonifera* and *F. brevipila* phenotypes a week after five fungicide applications. The plant treated with propiconazole fungicide showed darker green leaf color and slower growth (A, C). *Agrostis stolonifera* also showed wider leaves (A), while the water-treated plants showed normal growth and color (B, D).

### Transcriptome Assembly and Annotation

A total of 392,578,625 raw reads were generated to study plant-fungicide interaction; after adaptor removal and quality trimming, a total of 392,556,876 reads were retained. To study the different responses to pink snow mold inoculation in fungicide- and water-pretreated plants, a total of 970,893,276 reads were generated from Illumina sequencing. After adaptor removal and quality trimming, 919,321,188 reads were retained. All sequencing reads were used to build the *de novo* assembly that included 982,775 contigs with N50 of 1,047 bp. After combining *de novo* and reference transcriptome assemblies and removal of sequence with high similarity, we obtained a combined assembly with 734,009 contigs and N50 of 1,042 bp.

The BUSCO assessment showed 422 of the BUSCO genes were fragmented in the *de novo* assembly and the number was reduced to 185 in the combined assembly. Additionally, only 43 BUSCO transcripts were showing as missing in the hybrid assembly, while the *de novo* assembly potentially missed 113 genes (**Figure 2 A**). The annotation of combined assembly statistics was shown in **Figure 2 B** with 25,239 transcripts had annotation from all five databases. NCBI NR and UniRef had the most annotation for this combined assembly.

**Figure 2.**
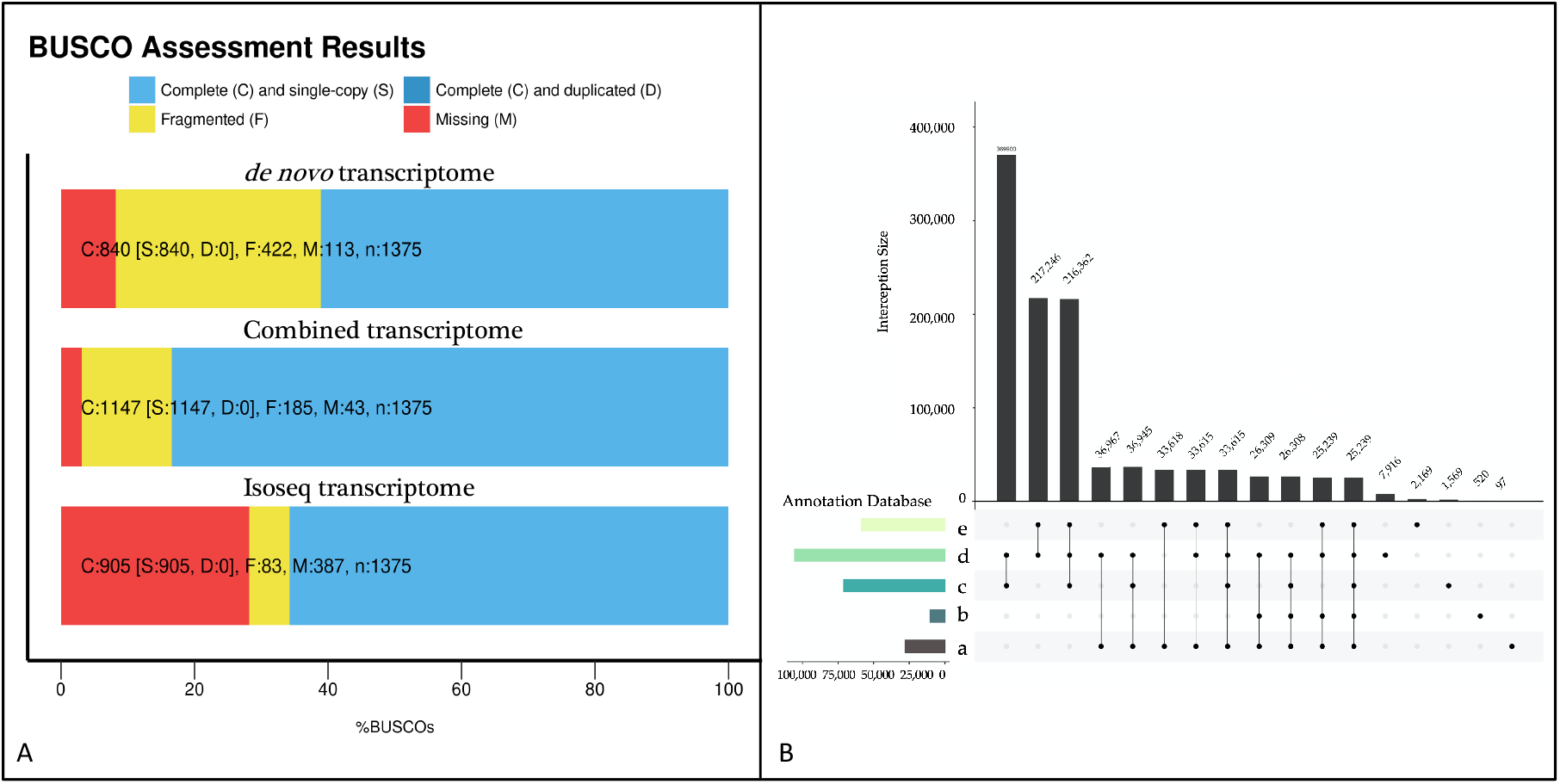
The transcriptome completeness evaluation (A) and transcriptome annotation using five protein databases (B). Protein databases in panel (B): a: Kyoto Encyclopedia of Genes and Genomes; b: Gene Ontology; c: NCBI NR Protein; d: UniRef90; e: SwisProt.

### Differentially Expressed Genes Analysis

RNA sequencing reads were mapped to the transcriptome reference using the combined assembly, isoseq only, and *de novo* assembly. The RNA sequencing reads mapping statistics are summarized in **Table 1**. The combined assembly had the least unmapped reads, followed by Isoseq reference. The *de novo* assembly had the most unmapped reads.

**Table 1.** RNA-Seq yield and reads mapping for the propiconazole study using Isoseq, *de novo*, and combine transcriptome, respectively. (PCZ: Propiconazole)

| Treatment | Replicates | Total reads | High quality reads | Reference | Mapped reads (%) | Uniquely mapped reads (%)<br>(MAPQ>3) |
| --- | --- | --- | --- | --- | --- | --- |
| PCZ | R1 | 62,266,322 | 62,262,686 | Isoseq | 64 | 57.1 |
|  |  |  |  | *de novo* | 48.6 | 44.7 |
|  |  |  |  | Combined | 72.4 | 66.8 |
|  | R2 | 72,353,963 | 72,349,738 | Isoseq | 62.7 | 57.1 |
|  |  |  |  | *de novo* | 48.1 | 44.2 |
|  |  |  |  | Combined | 71.9 | 66.2 |
|  | R3 | 65,990,410 | 65,986,554 | Isoseq | 65.5 | 59 |
|  |  |  |  | *de novo* | 45.9 | 42.9 |
|  |  |  |  | Combine | 72 | 66.9 |
| Water | R1 | 64,102,358 | 64,098,632 | Isoseq | 63.5 | 57.2 |
|  |  |  |  | *de novo* | 47.7 | 44 |
|  |  |  |  | Combined | 71.6 | 66.3 |
|  | R2 | 59,400,130 | 59,397,440 | Isoseq | 64.1 | 57.2 |
|  |  |  |  | *de novo* | 48.4 | 44.6 |
|  |  |  |  | Combined | 71.8 | 65.9 |
|  | R3 | 68,465,442 | 68,461,826 | Isoseq | 63.7 | 56.9 |
|  |  |  |  | *de novo* | 48.2 | 44.5 |
|  |  |  |  | Combined | 71.9 | 66.3 |

The RSEM mapping result using the combined assembly as the reference was used for differential gene expression analysis. At FDR 0.05 with a minimum fold change of 2.0, a total of 204 differential expressed genes (DEGs) were identified with 117 genes upregulated and 87 downregulated in propiconazole-treated plants (**Figure 3**). Based on the conserved domain, two of the 117 upregulated genes were identified as CYP450 genes.

**Figure 3.**
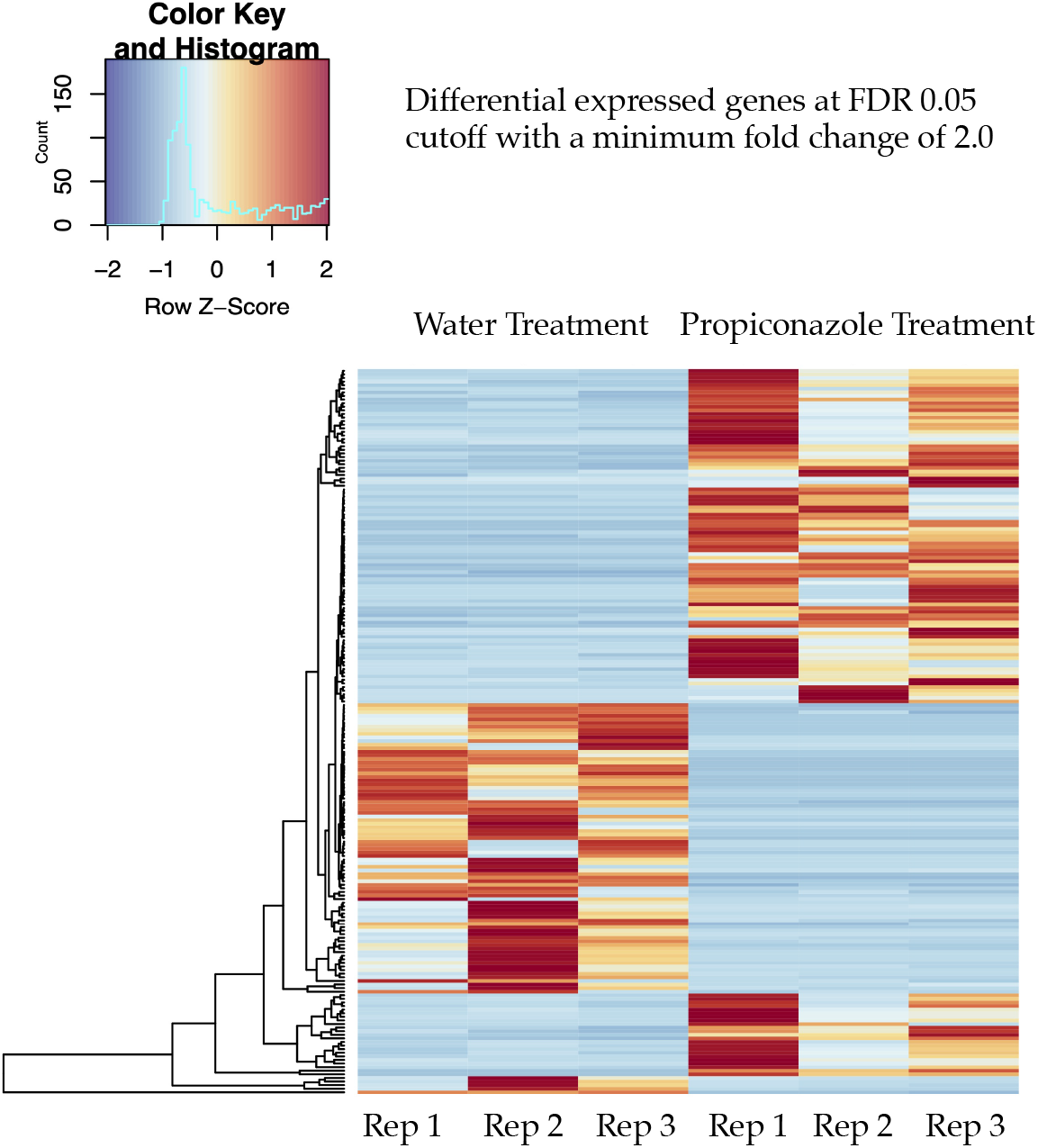
Differentially expressed genes between propiconazole-treated and water-treated *control* plants at FDR 0.05, minimum Log fold change of 2.0.

### CYP450 Related Differentially Expressed Genes

Transcript_34799 (logFC: −2.37) and Trinity_dn78694_c0_g4_i1(LogFC: −9.57) were identified as CYP450 genes. To provide a phylogenetic relationship between these two transcripts with known CYP450 genes, a total of 274 Arabidopsis and rice CYP genes were used as the reference. After sequence alignment and trimming, iqtree model testing suggested the JTT+F+I+G4 as the best model which was then used to reconstruct the phylogenetic tree. Transcript_34799 was a sister to rice CYP92A9 and Trinity_dn78694_c0_g4_i1 was close to the *Arabidopsis* CYP71A gene family. Both transcripts were nested the CYP71 Clan (**Figure 4**).

**Figure 4.**
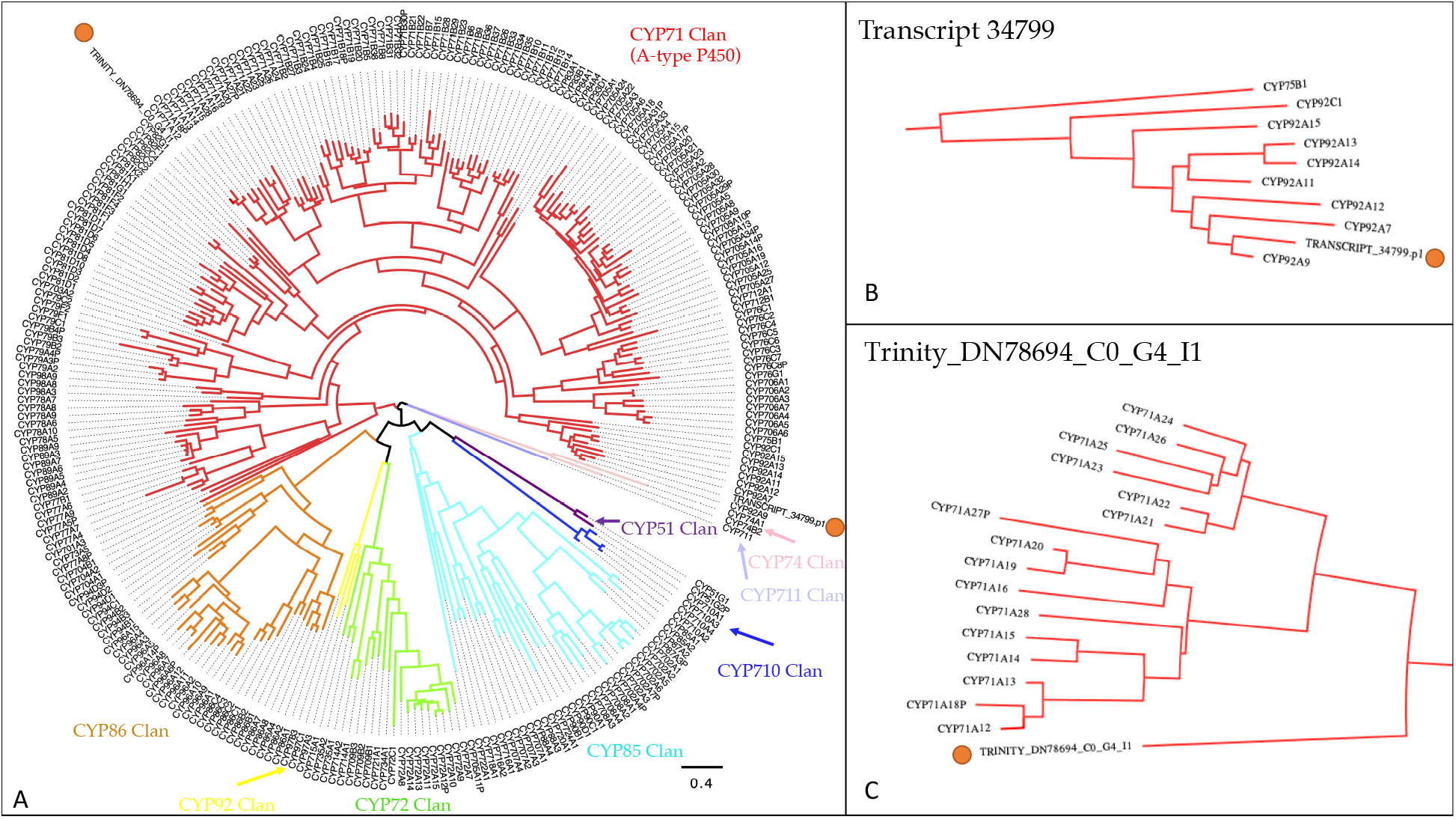
The phylogenetic relationship of the two P450 transcripts that were upregulated under the propiconazole treatment with Arabidopsis and rice P450 genes. Nine CYP clans were color coded, and the two transcripts from *F. brevipila* were highlighted in orange dots (A). The zoomed regions where the two transcripts were located (B, C). The transcript_34799 was a sister to rice CYP92A9 and Trinity_dn78694_c0_g4_i1 was close to the *Arabidopsis* CYP71A gene family.

The NCBI blastP results suggested Transcript_34799 had the best hit to CYP92A44-3 gene (GenBank: AER39773.1), with a max score of 772, 99.21% protein sequence similarity. The CYP92A44-3 was identified and cloned from *Festuca rubra* ssp. *fallax*, a close relative of *F. brevipila*. The second hit of this transcript was CYP71A genes in *Aegilops tauschii* ssp. *tauschii* (697, 90.50%), and followed by a flavonoid 3’-monooxygenase in *Hordeum vulgare*. Since the phylogenetic data suggested Transcript_34799 was a sister to the rice CYP92 gene family, the final annotation of this transcript in this study would be considered as CYP92A family gene.

The Trinity_dn78694_c0_g4_i1 transcript had highest sequence similarity to CYP450 71A1-like protein (XP_020192280.1) in *Aegilops tauschii* ssp. *tauschii*, and a CYP450 71A1 protein (GenBank: EMS50742.1) in *Triticum urartu*, which was within expectations of the phylogenetic data. The final annotation of this transcript in this study would be considered as a CYP71A family gene.

### Non-CYP450 Differentially Expressed Genes

The non-CYP450 DEGs were selected based on gene annotation using five databases, and further grouped by metabolic activities and transcriptional factor/signaling functions (**Figure 5**). For example, we observed upregulation of gibberellin 2-oxidase, cinnamoyl-CoA reductase, CBF protein, and multiple WRKY (5, 30, 41, 53) transcriptional factors induced in the propiconazole-treated plants while the expression of NADH ubiquinone oxidoreductase and cinnamyl alcohol dehydrogenase were downregulated. All the transcript ID and fold changes are summarized in **Table S2**.

**Figure 5.**
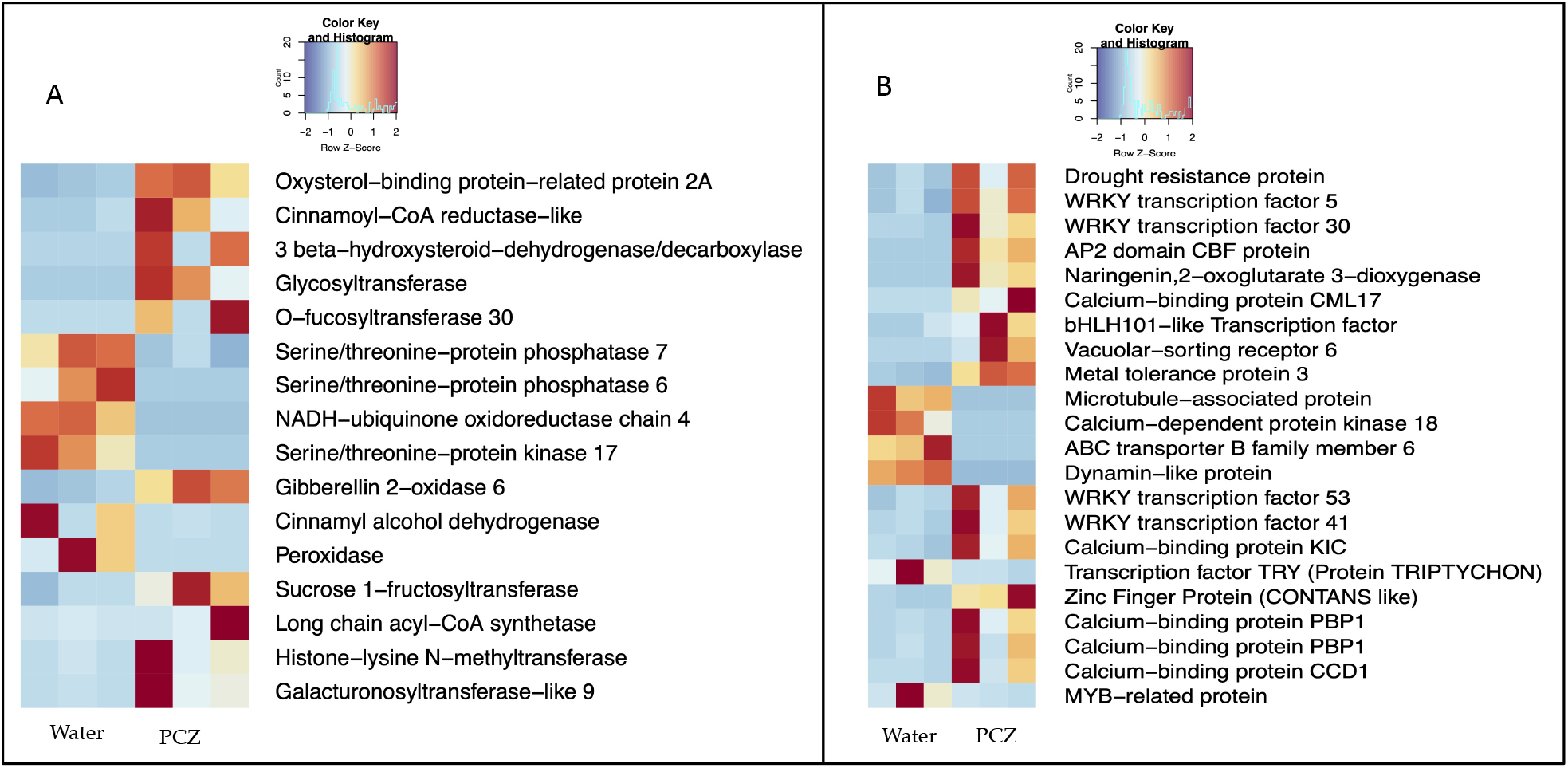
Subsets of enzymes (A) and transcriptional factors (B) affected by the propiconazole (PCZ) fungicide application in *F. brevipila*.

In addition to the transcriptional factors and metabolite-associated enzymes, such as photosystem II activity, several expansin-related genes activities were also down regulated in propiconazole-treated plants (**Table S2**).

### Metabolites Profile Change

The metabolite profile from the HILIC column run was analyzed with retention time set to 0.5-25 minutes. After quality control and CV-based filtering, the unsupervised metabolite clustering for positive (720 features) and negative (692 features) modes were shown in **Figure 6 (A, B)**, where principle components 1 (PC1) explained 37.8% and 16.4% variation in positive and negative mode, respectively. The initial quality control showed in one of the four subsamples from one of the water-treated plant was clustered with propiconazole treated plants in the C18 dataset. This was likely the result of sampling contamination. Since there were four subsamples from the same plant, this subsample was removed from the downstream analysis. The first pool sample was also removed in the HILIC column data analysis because it deviated from the center of the PCA.

**Figure 6.**
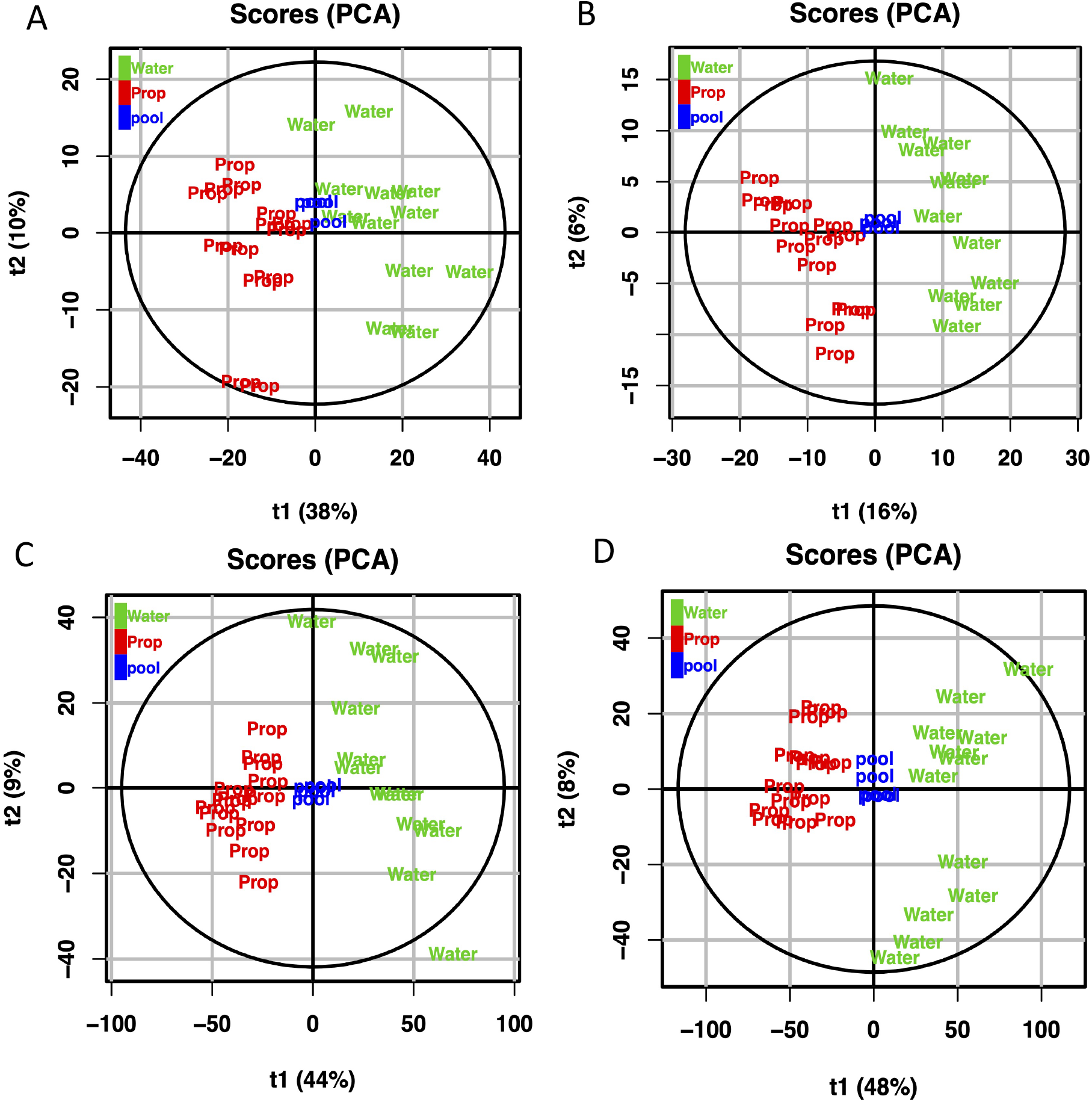
The multivariate analysis of metabolite features generated from HILIC column (A, B) and C18 column (C, D). The unsupervised metabolite clustering for positive and negative modes from both columns showed treatment effects on both primary and secondary metabolites profile.

The metabolite profile changes from the C18 column was analyzed using the same method except the retention time was subset to 1-15 minutes. The positive mode included 2,933 metabolite features, and the negative mode included 4,090 metabolite features. The unsupervised clustering for the metabolite profiles is shown in **Figure 6 (C, D)** with PC1 explaining 44.5% variation in positive mode, and 48.2% in negative mode.

The top 10 metabolites for each mode and column (a total of 40) predicted by accurate m/z mass are shown in **Table 2**. Because numerous metabolites could match with accurate mass data, more than one database hit was included for each feature.

**Table 2.**
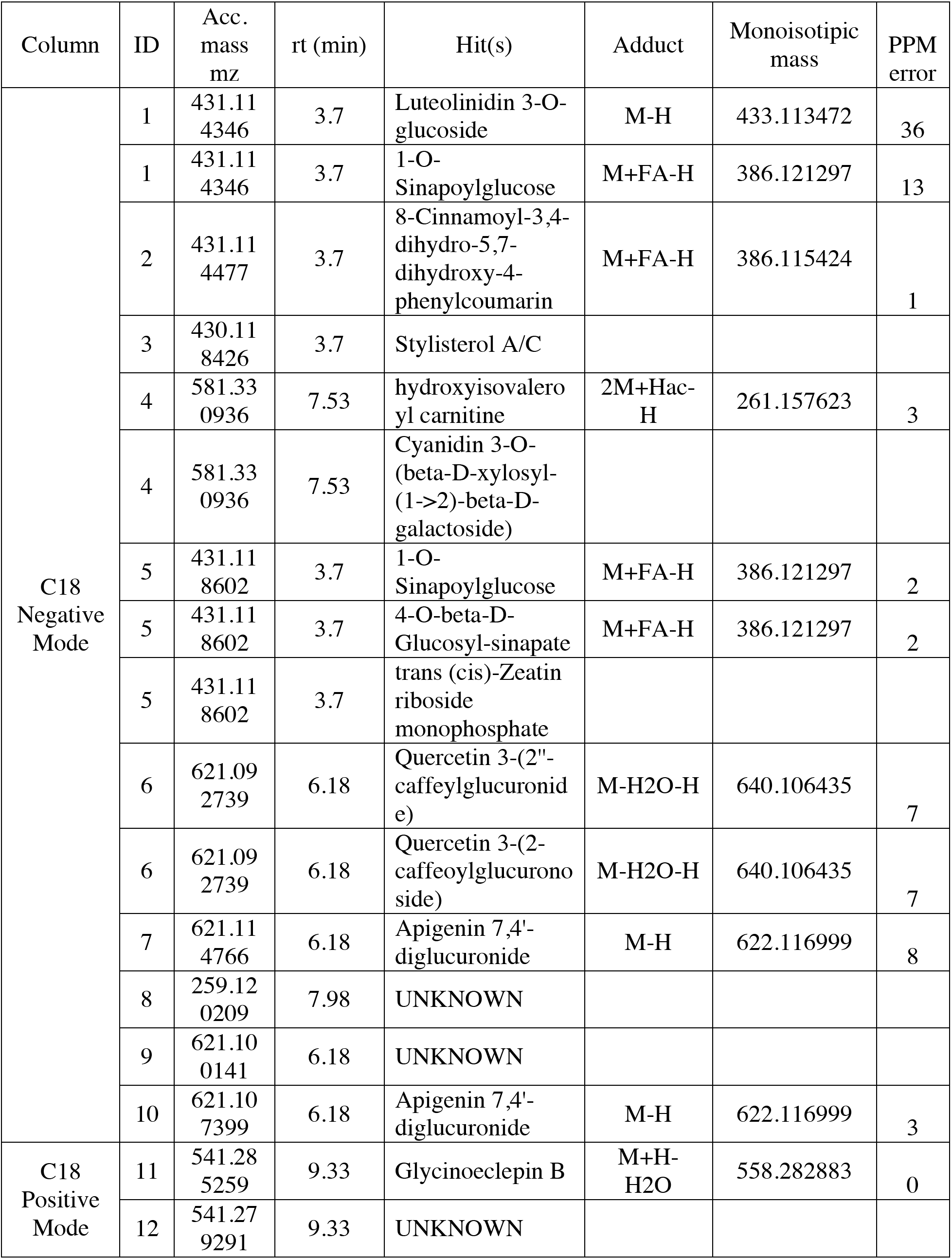
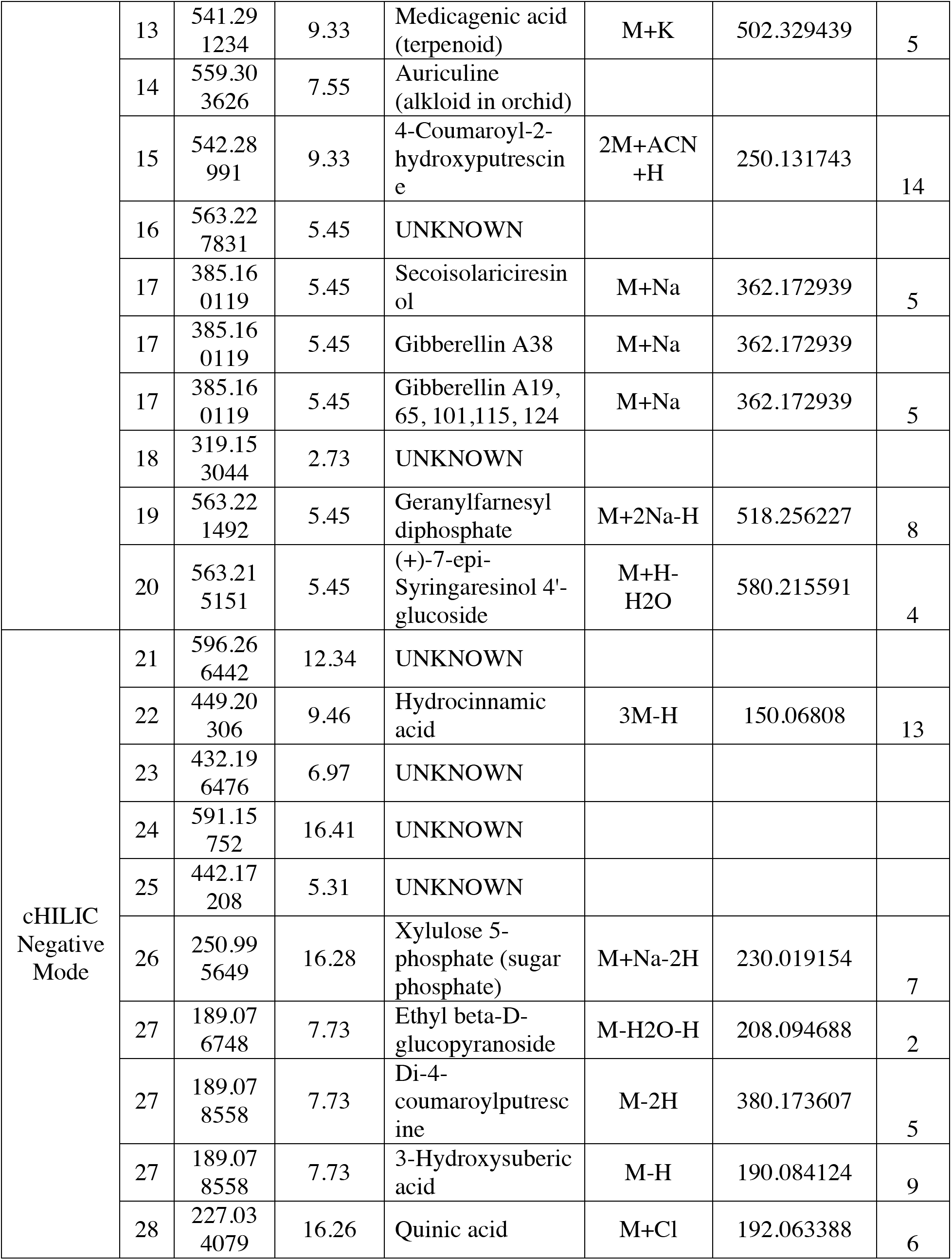
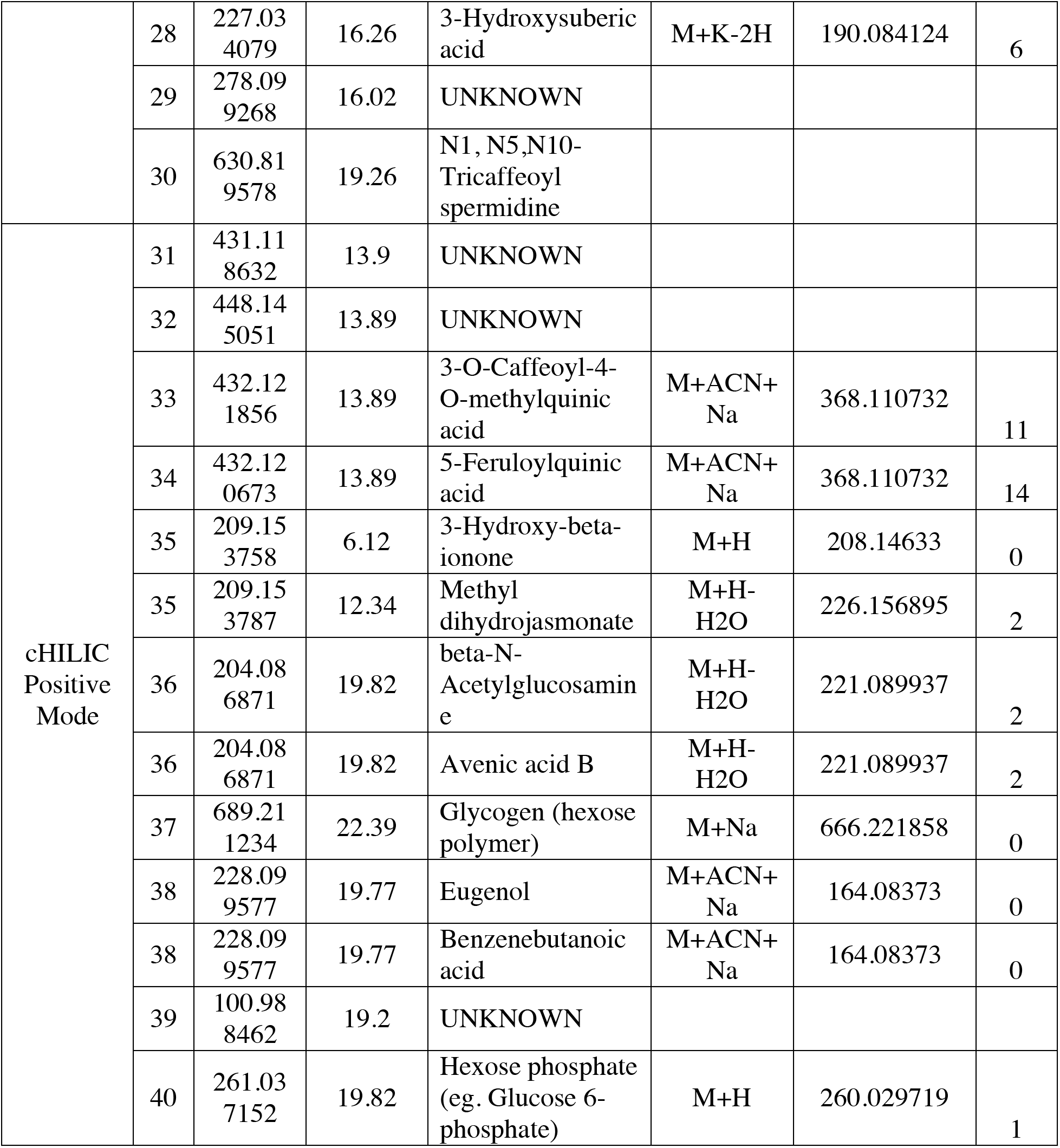
Accurate mass-based metabolite feature prediction using the human metabolome database and KEGG compound database.

In addition to the top 10 metabolites for each mode and column, we also identified a total of 93 candidate metabolites (C18: 92; cHILIC 1) that were unique only in propiconazole treated plants (**Table 2**)

### Propiconazole Pretreatment Increased the *M. nivale* Resistance of *F. brevipila*

Fourteen days post inoculation, WS plants had significant mycelium growth compared to FS plants (**Figure 7**). To compare gene expression difference between FS and WS one day post snow mold inoculation, RNA sequencing identified a total of 4,118 differentially expressed genes (DEGs) at FDR 0.05 with the minimum LogFC of 2.0. Filtered by NCBI NR protein database annotation, the 4,118 DEGs were split into *Microdochium* fungi-specific (1,730 DEGs) and plant-specific groups (1,871 DEGs). DEGs without NCBI NR annotation were excluded from further analysis. Heatmaps suggested a higher level of *Microdochium* activity in the WS group (**Figure S6 A**), and some *Microdochium* activity in the FS group; there was also a clear expression pattern separation in the plant-specific genes expression (**Figure S6 B**).

**Figure 7.**
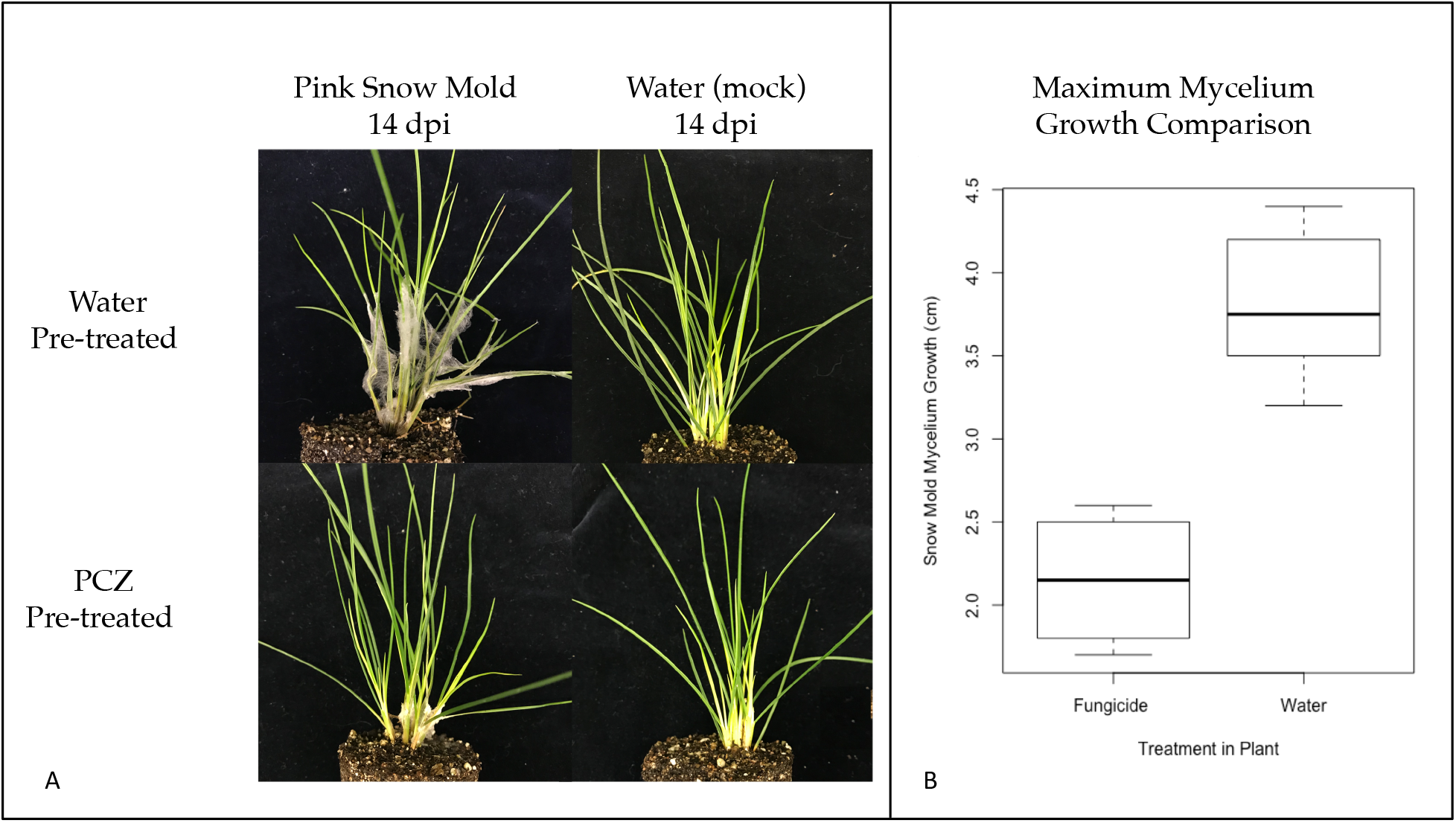
Pink snow mold inoculation showed WS plants had significantly more snow mold spread compared to other treatments. FS plants had some level of snow mold infection. PCZ: propiconazole; WS: water pretreated plant with snow mold; FS: fungicide pretreated plant with snow mold.

A total of 143 upregulated genes in the FS group had KEGG annotation and 821 upregulated genes in the WS group had KEGG annotation. KEGG mapper reconstruction suggested that specific pathway modules such as fatty acid and α-linolenic acid metabolism were upregulated in the WS group comparing to the FS group.

After inspecting NR-based annotation for each group using candidate genes information from previous literature, we found the ice recrystallization inhibition proteins (IRIPs), cold responsive proteins, and disease resistance proteins were upregulated in the FS group at significant levels, while chitinase and pathogenesis-related proteins had significantly higher expression in the WS group (**Figure S7, Table 3**). The conserved domain search of the six IRIPs showed four out of the six IRIPs contained at least one leucine-rich repeat receptor-like (LRR) protein kinase domain, and three contained ice binding domains (**Figure S8**). One identified IRIPs contained CAP (cysteine rich secretory proteins, antigen 5 and pathogenesis-related 1 protein domain).

**Table 3.** Cold and pathogen infection related DEGs between the FS and WS treatments.

| Transcript ID | NCBI NR Based Transcript Annotation | LogFC (FS/WS) | *p*-value |
| --- | --- | --- | --- |
| TRINITY_DN72353_C1_G2_I2 | Cold responsive protein | 2.60 | 2.0E-06 |
| TRINITY_DN69307_C0_G1_I7 | Glycine-rich cell wall structural protein | 3.34 | 2.0E-03 |
| TRINITY_DN88735_C0_G1_I2 | Cold-induced small protein | 3.79 | 1.5E-03 |
| TRINITY_DN74175_C0_G1_I10 | Disease resistance protein RGA2 | 7.77 | 1.9E-05 |
| TRINITY_DN74175_C0_G1_I3 | Disease resistance protein RGA2 | 2.66 | 1.3E-04 |
| TRANSCRIPT_12036 | Disease resistance protein RPM1-like | 2.18 | 1.9E-04 |
| TRINITY_DN81777_C0_G1_I12 | Disease resistance RPP13-like protein | 2.28 | 8.6E-04 |
| TRINITY_DN100146_C0_G1_I9 | Fructan 6-exohydrolase | 2.50 | 1.5E-04 |
| TRINITY_DN97131_C2_G5_I3 | Ice recrystallization inhibition protein 2 | 2.70 | 2.7E-03 |
| TRANSCRIPT_46772 | Ice recrystallization inhibition protein 5 | 2.11 | 1.5E-03 |
| TRANSCRIPT_48329 | Ice recrystallization inhibition protein 6 | 2.05 | 3.3E-04 |
| TRINITY_DN86392_C0_G9_I1 | Ice recrystallization inhibition protein 6 | 2.81 | 2.7E-04 |
| TRINITY_DN77946_C1_G1_I3 | Ice recrystallization inhibition protein IRI1 | 2.26 | 2.2E-03 |
| TRINITY_DN72600_C2_G3_I3 | Ice recrystallization inhibition protein IRI3 | 4.50 | 4.9E-04 |
| TRINITY_DN100575_C2_G2_I8 | Isoflavone reductase | 2.13 | 3.7E-04 |
| TRINITY_DN105886_C6_G1_I5 | Proline synthase | 8.02 | 4.0E-04 |
| TRANSCRIPT_3235 | Putative disease resistance protein | 2.16 | 6.5E-04 |
| TRANSCRIPT_53735 | Chitinase | -3.91 | 2.4E-03 |
| TRINITY_DN87228_C0_G3_I17 | Chitinase 1 | -3.82 | 1.0E-05 |
| TRINITY_DN80789_C0_G3_I4 | Chitinase 1 | -2.47 | 2.4E-03 |
| TRINITY_DN80789_C0_G3_I2 | Chitinase 1 | -3.53 | 1.6E-03 |
| TRANSCRIPT_54666 | Chitinase 1 | -3.13 | 1.4E-03 |
| TRANSCRIPT_50472 | Chitinase 8-like | -2.07 | 2.0E-05 |
| TRINITY_DN98644_C1_G1_I2 | Pathogen-related protein | -2.55 | 5.5E-05 |
| TRINITY_DN73403_C1_G1_I1 | Pathogen-related protein | -2.67 | 1.7E-03 |
| TRANSCRIPT_52665 | Pathogen-related protein-like | -3.08 | 3.9E-04 |
| TRINITY_DN91197_C0_G3_I1 | Pathogenesis-related protein 1 | -2.23 | 2.3E-04 |
| TRINITY_DN91197_C0_G2_I1 | Pathogenesis-related protein 1 | -2.23 | 1.0E-03 |
| TRANSCRIPT_49936 | Pathogenesis-related protein 1 | -2.80 | 2.8E-09 |
| TRANSCRIPT_47468 | Pathogenesis-related protein 1 | -6.88 | 9.6E-24 |
| TRINITY_DN91197_C0_G1_I4 | Pathogenesis-related protein 4 | -2.69 | 8.5E-05 |
| TRINITY_DN87228_C0_G3_I5 | Pathogenesis-related protein 4 | -3.47 | 8.5E-07 |
| TRINITY_DN87228_C0_G3_I4 | Pathogenesis-related protein 4 | -2.05 | 2.8E-04 |
| TRINITY_DN102652_C0_G1_I23 | Pathogenesis-related protein<br>PRB1-3 | -2.99 | 3.0E-06 |

## Discussion

Propiconazole is a triazole fungicide known as a demethylation inhibitor (DMI), that targets the C14-alpha demethylase enzyme from removing the C-14α-methyl group from lanosterol to stop ergosterol production in fungi (Peyton, Gallagher et al. 2015). PCZ is a known regulator of plant growth (Fletcher, Gilley et al. 2000) and has been shown to enhance root development of redroot pigweed (*Amaranthus retroflexus*) (Hanson, Mallory-Smith et al. 2003). PCZ treatment has also been shown to reduce salt stress in Kentucky bluegrass (*Poa pratensis* L.) (Nabati, Schmidt et al. 1994), speed up the recovery of Kentucky bluegrass sod from heat injury (Zhang, Ervin et al. 2003), and provide a greater turf quality in creeping bentgrass (Ervin, Zhang et al. 2004) by increasing superoxide dismutase (SOD) activity. In addition, PCZ was found to be an inhibitor for brassinosteroid biosynthesis in *Arabidopsis* and maize (Sekimata, Han et al. 2002, Hartwig, Corvalan et al. 2012, Oh, Matsumoto et al. 2016) and also showed an inhibition effect on obtusifoliol 14R-demethylase activity (Burden, Cooke et al. 1989, Gilley and Fletcher 1997).

Major brassinosteroid biosynthetic genes include but are not limited to, cytochrome P450s 85A, 90B, 90C, 90D and 92A (Nomura and Bishop 2006). Studies have shown BR inhibitors, such as brassinazole and triadimefon, bind to CYP90B, C and D to suppress BR biosynthesis (Asami, Mizutani et al. 2003). In our study, after five propiconazole applications, the positive control creeping bentgrass samples used in this study showed darker leaf color, wider leaves, and stunted growth as expected. Because the leaves of hard fescue are narrow, we did not measure leaf size change. We observed the dwarf phenotype with darker green leaf colors. Through transcriptome analysis, we identified two CYP 450 (CYP92A and CYP71A) that were upregulated in PCZ-treated plants. CYP92A was found to convert typhasterol to castasterone both early and late in the C6 brassinosteroid biosynthesis pathway and was believed to function as BR C-2 hydroxylase (Nomura and Bishop 2006). The application of P450 inhibitors [Ancymidol and 1-amin-obenzotriazole (ABT)] on Chewings fescue (*Festuca rubra* ssp. *fallax*), a close relative of *F. brevipila*, also showed the upregulation of CYP92A gene (Huang, Rehak et al. 2012).

Our finding agrees with previous studies that in BR-deficient plants, genes committed to BR biosynthesis were upregulated at the transcription level (Noguchi, Fujioka et al. 2000). In our case, we did not observe differential expression of CYP90 and CYP85 genes. One possible reason is that CYP90A1 and CYP85A2 have a circadian rhythm of gene expression and are not affected by the feedback regulation (Bancos, Szatmári et al. 2006), or simplify because the variation induced by leaf differs with age.

The CYP71A gene family is thought to have highly specific functions across and within individual species (Hamberger and Bak 2013). For example, the CYP71A1 gene was suggested to play a role in oxidation of monoterpenoids in avocado fruit (Bozak, Yu et al. 1990), while CYP71A13 was found to be involved in the camalexin biosynthesis (Nafisi, Goregaoker et al. 2007). In our study, the protein identified as CYP71A1-like did not cluster close to a specific CYP71A gene in *Arabidopsis*; thus, its function remains unknown.

Sekimata et al. (2012) suggested that PCZ affects the BR biosynthetic pathway through a BR independent pathway, while Asami et al. (2003) suggested triadimefon affects both the GA and BR pathways. In our study, we observed the up-regulation of giberellin 2-oxidiase 6 in PCZ-treated plants. The activiation of giberellin 2-oxidiase 6 was found to deactivate active giberellin in rice and created a semi-dwarf phenotype (Huang, Tang et al. 2010). Interestingly, GA and BR interact via DELLA proteins to regulate plant growth and development (Li, Wang et al. 2012), and the dwarf phenotype we observed in this study could be the combination of BR and GA pathway effects. Future study is needed to address this question.

Through transcriptome analysis, we also observed that multiple plant signaling, and defense pathways were upregulated by PCZ treatment. The upregulation of WRKY 30 has been associated with plant salt resistance by regulating reactive oxygen species-scavenging in grape (Zhu, Hou et al. 2019), and increased endogenous jasmonic acid accumulation and PR gene expression in rice (Peng, Hu et al. 2012). WRKY5 was found to be related to SA signaling and induced by elicitors (Cormack, Eulgem et al. 2002, Schilirò, Ferrara et al. 2012). The CBF family is usually regulated by cold temperatures (Gilmour, Zarka et al. 1998, Medina, Bargues et al. 1999). Interestingly, we also observed the upregulation of the AP2 domain CBF protein. These results suggest the fungicide potentially functions similar to a stress stimulus, such as cold stress.

We observed some gene expression variations within biological repeats that might be due to the leaf age variations during the sampling. Metabolomics feature (C18 and HILIC) scatter plots using unsupervised clustering showed separation between propiconazole and water-treated plants, suggesting a treatment effect on *F. brevipila*. Of the forty features driving the feature clustering of C18 and cHILIC metabolite profiles, metabolites such as methyl dihydrojasmonate, gibberellin acid isomers, secoisolariciresinol (lignan, a type of phenylpropanoid compound), and 4-O-beta-D-glucosyl-sinapate (a lignin monomer with a hexose) are particularly interesting because transcriptome data suggested potential changes in GA signaling pathway (GA 2-oxidase), jasmonic acid accumulation (WRKY30), and lignin biosynthesis [cinnamoyl-CoA reductase, cinnamyl alcohol dehydrogenase (Kawasaki, Koita et al. 2006, Ma 2010)]. However, the accurate mass search returned with multiple putative metabolites. It is difficult to confirm the metabolite features based on information available. Future research will be done to using standard compounds, tandem MS, or even nuclear magnetic resonance (NMR) to characterize and confirm the predicted metabolites from this dataset.

We found that hard fescue plants, 6 months after propiconazole treatment, showed better resistance to snow mold than plants treated with water. Pink snow mold resistance is associated with the winter hardiness of the plant. In winter rye, chitinase, thaumatin-like protein (GLP), and endo-β-1,3-glucanase (GLPs) were found to confer snow mold disease resistance (Hiilovaara-Teijo, Hannukkala et al. 1999, Kuwabara, Takezawa et al. 2002). Multiple reports suggested the cold hardiness-related traits such as cold-induced PR, GLP proteins have antifungal activities (Bertrand, Castonguay et al. 2011, Kovi, Abdelhalim et al. 2016). Cytological analysis on wheat suggested that cold hardened wheat delayed the infection of the snow mold penetration and showed better disease resistance compared to the non-harden plants (Dubas, Golebiowska et al. 2011). The modification of cell wall components was also found to be associated with the snow mold resistance in perennial ryegrass (Kovi, Abdelhalim et al. 2016). In our study, when comparing the fungicide- and water-pretreated plants, transcriptome data showed more pink snow mold gene activity in plants that were pretreated with water. This suggests that the pathogen had a faster penetration or growth on water-pretreated plants in the first 24 hours. When comparing the PR gene expression between WS and WC groups, we observed the significant upregulation of PR proteins in WS, suggesting the expression of PR proteins were likely triggered by the pathogen and not the cold temperature. We observed multiple chitinases, pathogenesis-related (PR) protein, and GLP protein that were upregulated upon infection in WS plants, suggesting the activation of a pathway by the plant similar to a snow mold pathogen defense pathway.

Interestingly, we identified a set of highly up-regulated ice recrystallization inhibition proteins (IRIP), cold responsive proteins, and some disease resistance proteins expression in FS plants compared to the WS group. The primary function of the IRIPs is to inhibit ice recrystallization and minimize the physical damage caused by larger ice crystals, thereby enhancing plant freezing tolerance (Griffith, Antikainen et al. 1997, Tremblay, Ouellet et al. 2005, John, Polotnianka et al. 2009, Zhang, Fei et al. 2010). Additionly, the IRIPs have the LRR domain to the N-terminal protion and studies have suggested the pathogenesis-related function from the IRIP (Hon, Griffith et al. 1995, Yeh, Moffatt et al. 2000, Tremblay, Ouellet et al. 2005). The expression of IRIPs and disease resistance proteins are likely the result of a combination of pink snow mold and cool temperature stress. It is still unknown if the IRIPs produced in the propiconazole-pretreated plants have antifungal activity and the function of the disease resistance protein; future research such as *in vitro* expression of these genes in *E. coli* will provide evidence for their potential function beyond ice recryslization inhibtion function. The disease resistance protein RGA2 was found to restrict the pathogen growth (Song, Bradeen et al. 2003). One possible hypothesis based on these results is that the plants gained pink snow mold resistance as a result of the fast expression of cold responsivie protein, disease resistance protein, and IRIPs expression.

Plants used in the snow mold study were treated with large amounts of fungicide six months prior to snow mold inoculation. The half-lives of propiconazole in organic-rich soil is 20 days (Thorstensen and Lode 2001). To avoid the fungicide residual in the soil or plant tissue, we used newly developed tillers and transplanted to fresh soil twice to conduct the experiment six months post fungicide treatment. Considering the additional frequent watering, we do not expect that our observations are a result of fungicide residual interference. Both FS and WS plants have an identical genetic background; thus, the observed difference to cold and snow mold infection likely resulted from the rapid use of propiconazole. During the fungicide spray, the TRINITY_DN72600_C2_G3_I3 transcript, which contained ice binding and multiple LRR domain, had higher expression but not at a significant level due to one outlier in the control group (LogFC 0.61, p-value 0.36; average transcript count: 214 to 133). It has been shown non-low-temperature stresses such as drought can also trigger antifreeze activity-related gene expression (Yu and Griffith 2001). It is possible that the propiconazole treatment induced a similar effect so when the plants are exposed to a similar condition, the plants are able to respond faster. Since we observed the potential propiconazole effect on lignin biosynthesis one-week post fungicide application, future cytogenetics work will be done to see if there is a change in cell wall structure.

## Conclusion

Fungicide applications are widely used in turfgrass research to create fungal endophyte-free plant populations to study the effect of fungal endophytes. We found that fungicide applications at rates and frequencies typically used in endophyte research affected the brassinosteroid and gibberellins pathways of plants. The long-term effect of the use of fungicide induced stronger disease resistance to *Microdochium nivale*, likely through the expression of cold responsive genes, disease resistance proteins, and anti-freezing proteins. For this reason, researchers should think carefully about fungicide use in fungal endophyte – grass host research experiments and results from these studies should be taken with caution. In addition, the cold-responsive, disease-resistance, and anti-freezing proteins identified in this study potentially provide plant breeders the new methods for developing turfgrass cultivars for better pink snow resistance.

## Acknowledgments

The authors would like to thank Minnesota Supercomputing Institute for providing computational resources. This research is funded by the National Institute of Food and Agriculture, USDA, Specialty Crop Research Initiative under award number 2017-51181-27222

