## Supplementary material for "Evaluating the Effects of Propiconazole on Hard Fescue (*Festuca brevipila*) via RNA sequencing and Liquid Chromatography–Mass Spectrometry": supplimental figures and table: Supplemental_Figures_tables1_2.docx

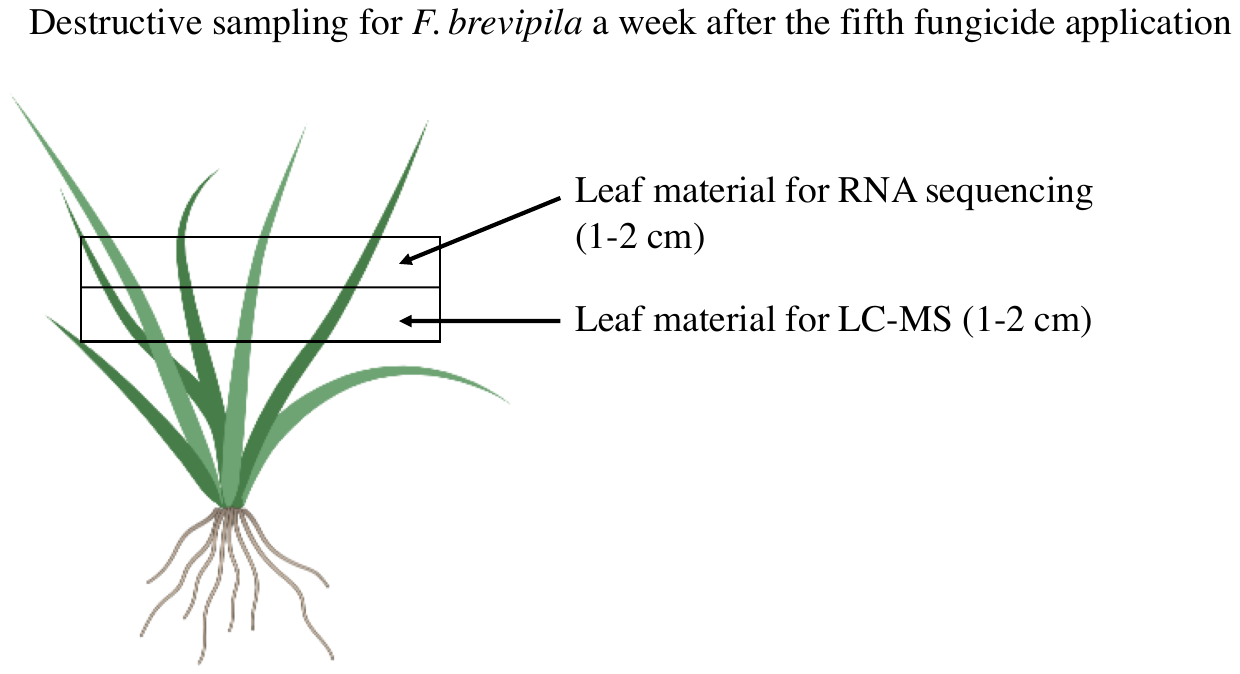

**Figure S1**. Destructive sampling of F. brevipila a week after the fifth fungicide application for transcriptome and metabolomics studies. For transcriptome sequencing, leaf samples were taken 1-2 cm from the leaf tip.

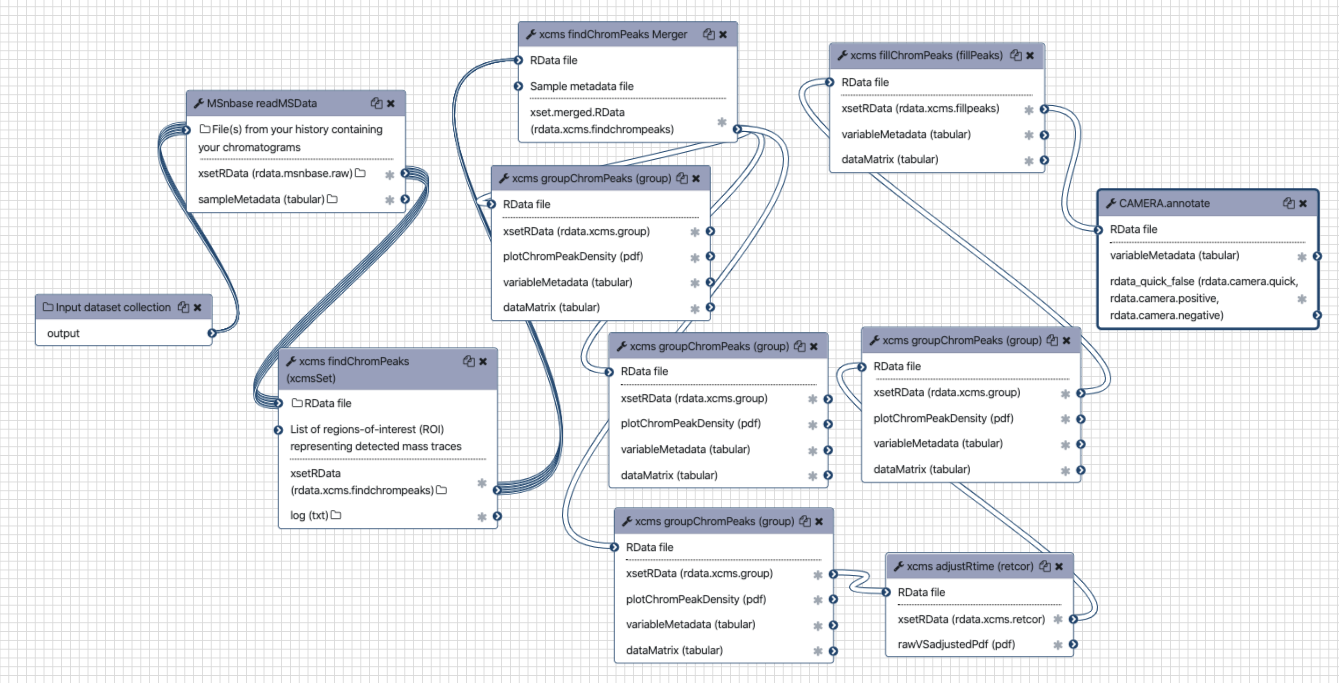

#### **Figure S2**. Galaxy metabolites data preprocessing workflow.

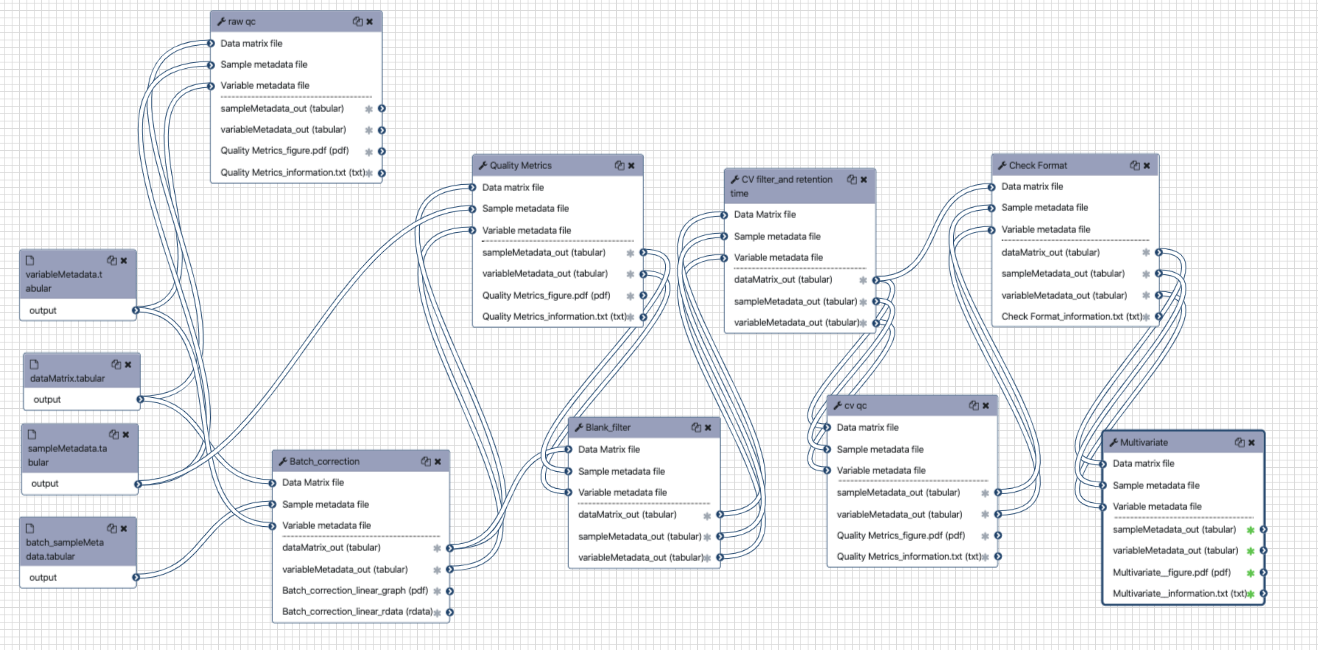

#### **Figure S3**. Galaxy metabolites data postprocessing workflow.

**Figure S4**. Workflow in preparation of plants for snow mold inoculation. Throughout the process, plants were transplanted twice into new soil. The snow mold inoculation was carried out six months after the last fungicide treatment.

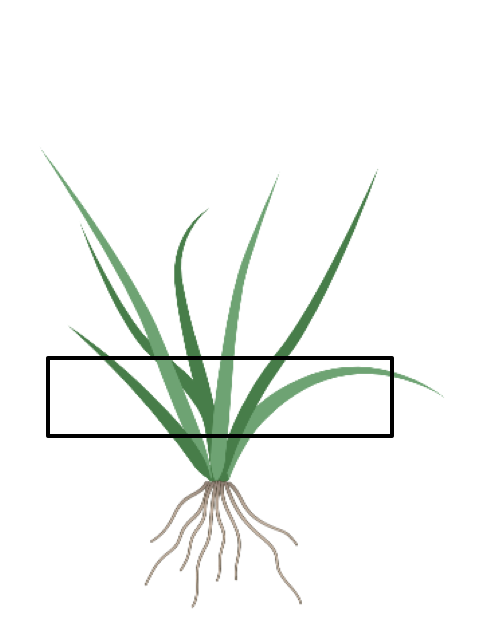

#### **Figure S5**. Figure illustration of plant tissues sampled for RNA sequencing in the long-term effect study.

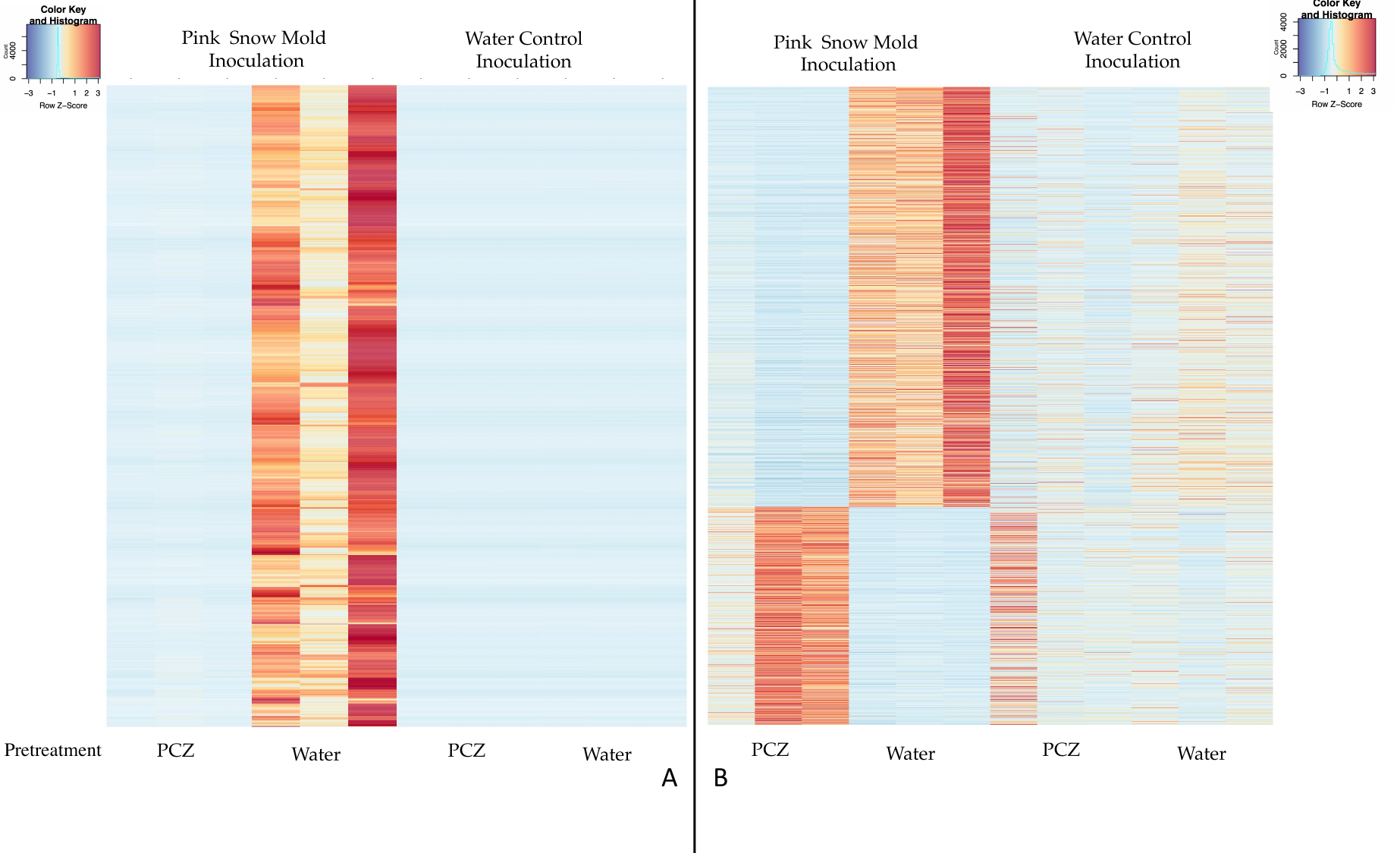

#### **Figure S6**. Differentially expressed Microdochium-related genes (A), and plant-related genes (B). Propiconazole (PCZ)

#### **Figure S7**. Cold and pathogen infection related differentially expressed genes between the fungicide pretreated plants with and without snow mold inoculation vs water pretreated plants with and without snow mold inoculation.

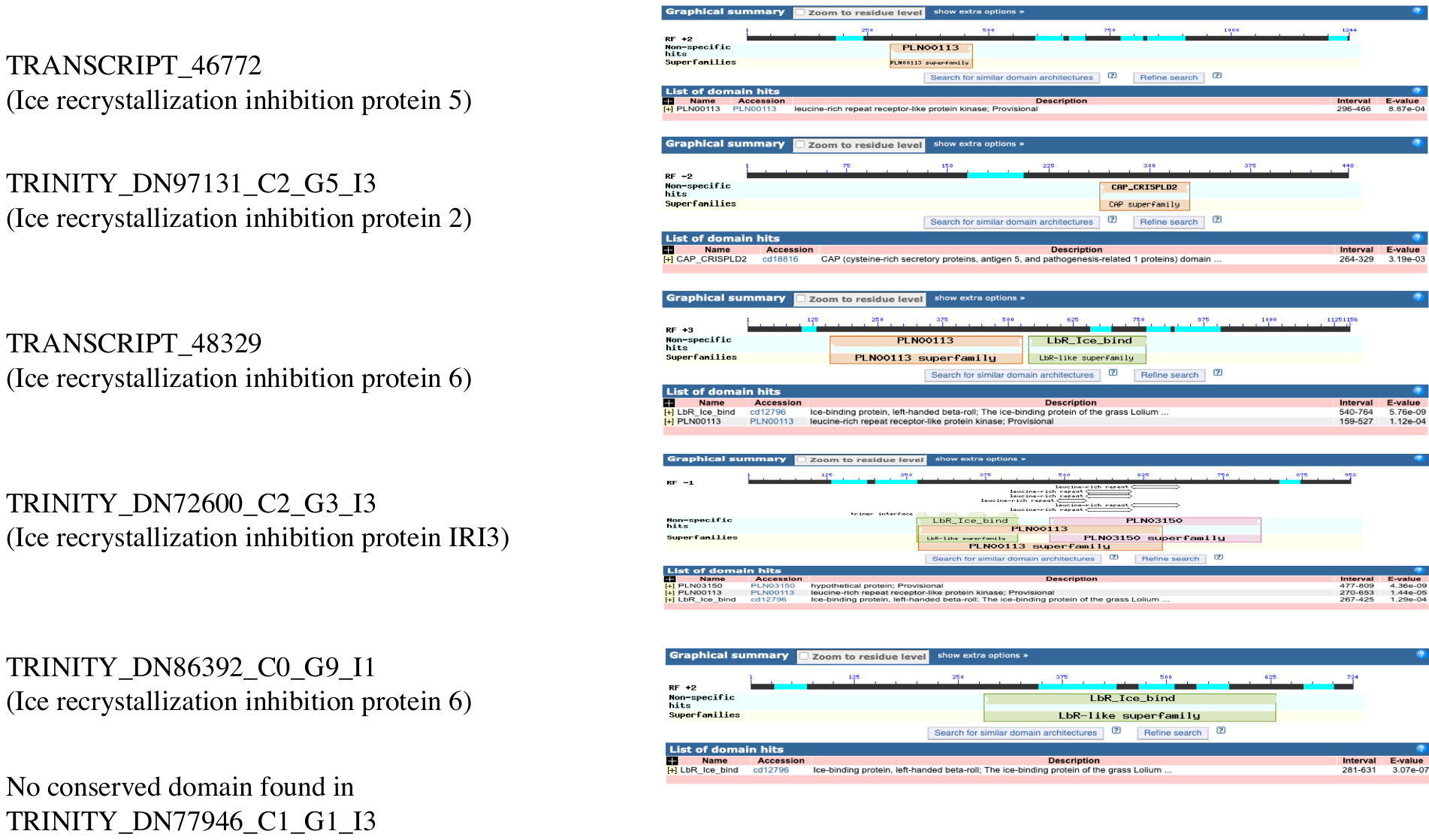

#### **Figure S8**. Conserved domain search in upregulated IRIPs via NCBI blastp

#### **Table S1.** Eight CYP92 genes downloaded from the rice genome database for the phylogenetic tree reconstruction.

| Gene ID | Gene Name |
| --- | --- |
| LOC_Os02g29960.1 | CPY92A15 |
| LOC_Os03g44740.1 | CYP92C1 |
| LOC_Os09g26980.1 | CYP92A7 |
| LOC_Os09g08920.1 | CYP92A13 |
| LOC_Os09g08990.1 | CYP92A14 |
| LOC_Os09g26940.1 | CYP92A11 |
| LOC_Os09g26960.1 | CYP92A9 |
| LOC_Os08g35510.1 | CYP92A12 |

#### **Table S2**. Selected downregulated genes in propiconazole-treated plants

| TRANSCRIPT_17444 | AEV91173.1 | 2.67 | MYB-related protein |
| --- | --- | --- | --- |
| TRANSCRIPT_45477 | AAW88315.1 | 4.70 | Expansin EXPA11 |
| TRANSCRIPT_45494 | XP_010228647.1 | 3.61 | peroxidase |
| TRANSCRIPT_46420 | CAC06433.1 | 3.23 | Expansin |
| TRANSCRIPT_46942 | CAC40806.1 | 4.69 | Beta expansin B3 |
| TRANSCRIPT_50937 | CAC40805.1 | 3.81 | Beta expansin B2 |
| TRANSCRIPT_56300 | AAT67050.1 | 3.10 | pathogenesis-related protein 4 |
| TRINITY_DN100223_C1_G3_I3 | AJK93406.1 | 6.23 | Cinnamyl alcohol dehydrogenase |
| TRINITY_DN100821_C1_G2_I9 | XP_020168398.1 | 9.37 | dynamin-like protein ARC5 |
| TRINITY_DN102104_C1_G1_I1 | EMS62391.1 | 9.17 | Splicing factor U2af large subunit A |
| TRINITY_DN104757_C2_G2_I1 | NP_037635.1 | 10.83 | RNA-dependent RNA polymerase |
| TRINITY_DN105176_C4_G2_I3 | XP_020172503.1 | 8.47 | microtubule-associated protein futsch isoform X1 |
| TRINITY_DN105573_C2_G1_I2 | XP_020186500.1 | 8.70 | putative glucuronosyltransferase PGSIP8 |
| TRINITY_DN75902_C1_G1_I6 | XP_010230204.1 | 8.37 | DNA (cytosine-5)-methyltransferase CMT1-like |
| TRINITY_DN76758_C2_G2_I10 | ACB45302.1 | 8.56 | expansin EXPA11 |
| TRINITY_DN79436_C4_G1_I1 | XP_020201563.1 | 2.80 | photosystem II |
| TRINITY_DN82258_C5_G1_I15 | XP_020162795.1 | 8.05 | Transportin MOS14 |
| TRINITY_DN82950_C1_G1_I7 | XP_024315534.1 | 8.00 | serine/threonine-protein phosphatase 6 |
| TRINITY_DN91298_C0_G1_I8 | XP_010228785.2 | 8.38 | ABC transporter B family member 6 |
| TRINITY_DN92704_C1_G2_I5 | XP_003571639.1 | 8.09 | exonuclease V, chloroplastic |
| TRINITY_DN97393_C1_G1_I3 | BAD87988.1 | 2.15 | putative beta-1,3-glucanase precursor |
| TRINITY_DN97393_C1_G2_I6 | AAU11328.1 | 3.92 | beta-1,3-glucanase 2a |
| TRINITY_DN98820_C0_G1_I8 | XP_020173114.1 | 7.86 | pentatricopeptide |
| TRANSCRIPT_23832 | XP_004975013.1 | -2.29 | alpha-humulene synthase |
| TRANSCRIPT_24929 | Q9FSV7.1 | -2.10 | sucrose 1-fructosyltransferase |
| TRANSCRIPT_33779 | XP_020170118.1 | -4.37 | ALP1-like protein |
| TRANSCRIPT_34799 | AER39773.1 | -2.37 | CYP92A44-3 |
| TRANSCRIPT_43011 | ACD80366.1 | -2.19 | WRKY5 transcription factor |
| TRANSCRIPT_44511 | ASU89566.1 | -2.11 | drought resistance |
| TRANSCRIPT_46869 | XP_020196033.1 | -2.31 | RING-H2 finger protein ATL3-like |
| TRANSCRIPT_47987 | XP_020174323.1 | -2.29 | histone-lysine N  methyltransferase 2D-like |
| TRANSCRIPT_53911 | XP_020171700.1 | -3.44 | adenomatous polyposis |
| TRANSCRIPT_55428 | XP_020198520.1 | -3.87 | heavy metal-associated isoprenylated plant protein 47-like |
| TRANSCRIPT_55823 | XP_003563436.1 | -3.59 | calcium-binding protein PBP1 |
| TRANSCRIPT_56908 | AAF86307.1 | -4.05 | EF-hand Ca2+ -binding protein CCD1 |
| TRINITY_DN104570_C1_G1_I7 | EMS68053.1 | -7.80 | Eukaryotic |
| TRINITY_DN105516_C3_G2_I17 | XP_003573998.1 | -9.81 | fluG protein |
| TRINITY_DN105968_C8_G1_I3 | XP_020179664.1 | -9.84 | RNA uridylyltransferase 1-like |
| TRINITY_DN71649_C4_G1_I1 | XP_020162934.1 | -7.68 | CO(2)-response secreted protease-like |
| TRINITY_DN72232_C0_G2_I2 | XP_020163886.1 | -6.29 | zinc finger protein CONSTANS-LIKE 1-like isoform X2 |
| TRINITY_DN73568_C4_G1_I3 | XP_020174579.1 | -3.00 | downy mildew resistance protein |
| TRINITY_DN73598_C0_G1_I4 | XP_003577532.1 | -9.68 | RNA polymerase |
| TRINITY_DN75106_C1_G2_I9 | AQK83574.1 | -8.15 | NAC domain-containing |
| TRINITY_DN76083_C0_G1_I4 | AFR67777.1 | -8.22 | AP2 domain CBF protein |
| TRINITY_DN76820_C2_G1_I3 | EMS67097.1 | -8.39 | putative 6-phosphogluconolactonase |
| TRINITY_DN78461_C5_G1_I12 | XP_020103799.1 | -2.52 | protein ROOT HAIR DEFECTIVE |
| TRINITY_DN79264_C0_G1_I7 | EMS56979.1 | -7.94 | Protein SPA1-RELATED 3 |
| TRINITY_DN82257_C0_G1_I13 | XP_003557393.1 | -8.56 | protein glycosyltransferase |
| TRINITY_DN84047_C5_G2_I3 | XP_003557748.1 | -8.15 | 3beta-hydroxysteroid-dehydrogenase/decarboxylase |
| TRINITY_DN92959_C3_G2_I9 | XP_020182066.1 | -7.43 | ras-related protein RABA1f-like isoform X2 |
| TRINITY_DN93007_C0_G3_I5 | EMS57783.1 | -9.02 | Naringenin,2-oxoglutarate |
| TRINITY_DN93029_C0_G1_I8 | XP_020193397.1 | -3.48 | transcription factor bHLH101-like |
| TRINITY_DN93083_C0_G1_I13 | XP_020164890.1 | -11.23 | bromodomain-containing protein |
| TRINITY_DN95724_C1_G1_I12 | XP_020177953.1 | -3.80 | long chain acyl-CoA synthetase |
| TRINITY_DN98193_C2_G1_I3 | XP_003570531.1 | -2.44 | metal tolerance |
| TRINITY_DN98261_C0_G2_I5 | XP_020188012.1 | -8.69 | AT-hook motif nuclear-localized |
| TRINITY_DN98980_C0_G1_I9 | XP_003558564.1 | -8.07 | peptidyl-prolyl cis-trans |
| TRINITY_DN99735_C0_G1_I2 | XP_003563336.1 | -9.14 | vacuolar-sorting receptor 6 |
