## Supplementary material for "Evaluating the Effects of Propiconazole on Hard Fescue (*Festuca brevipila*) via RNA sequencing and Liquid Chromatography–Mass Spectrometry": supplimental figures and table: Table S3_upload.docx

#### **Table S3**. Unique metabolite features that were only present in propiconazole-treated plants.

| Column | variableMetadata | mz | Rt (min) | isotopes | adduct | PCA_XLOAD-p1 | PCA_XLOAD-p2 |
| --- | --- | --- | --- | --- | --- | --- | --- |
| cHILIC Positive | M483T454 | 483.184001 | 7.56 | [268][M]+ |  |  |  |
| C18 Positive | M571T432 | 571.347308 | 7.20 |  |  | -0.0139596 | -0.0188306 |
|  | M593T441_1 | 593.355206 | 7.35 |  |  | -0.0114425 | -0.0174622 |
|  | M593T441_2 | 593.362072 | 7.35 |  |  | -0.0115762 | -0.0175756 |
|  | M593T440 | 593.368887 | 7.34 |  |  | -0.0123899 | -0.0168769 |
|  | M608T619_2 | 608.119962 | 10.32 |  | [M+H]+ 607.11 | -0.0208268 | -0.0206162 |
|  | M608T640_1 | 608.120887 | 10.66 |  |  | -0.0171249 | -0.0158836 |
|  | M608T619_3 | 608.127042 | 10.32 |  | [M+H]+ 607.11 | -0.0204855 | -0.0222937 |
|  | M608T640_2 | 608.128 | 10.66 |  |  | -0.0173484 | -0.0158309 |
|  | M608T619_4 | 608.134148 | 10.32 | [678][M]+ | [M+K]+ 569.175 [M+Na]+ 585.148 [M+H]+ 607.13 | -0.020523 | -0.0222508 |
|  | M608T619_5 | 608.141176 | 10.31 |  | [M+K]+ 569.175 [M+Na]+ 585.148 [M+H]+ 607.13 | -0.0204305 | -0.0166609 |
|  | M615T449_1 | 615.37466 | 7.49 |  |  | -0.0130924 | -0.0184822 |
|  | M615T449_2 | 615.381948 | 7.49 |  |  | -0.0125187 | -0.0186049 |
|  | M616T449 | 615.874948 | 7.49 | [690][M+1]2+ |  | -0.010771 | -0.019124 |
|  | M637T456_1 | 637.375532 | 7.60 | [727][M]2+ |  | -0.0113154 | -0.0165058 |
|  | M637T456_2 | 637.383195 | 7.61 |  |  | -0.0135044 | -0.0179125 |
|  | M637T456_3 | 637.390698 | 7.61 | [728][M]2+ |  | -0.0118337 | -0.0167707 |
|  | M637T457 | 637.398435 | 7.61 | [729][M]2+ |  | -0.0136284 | -0.0175941 |
|  | M638T456_2 | 637.88751 | 7.60 | [728][M+1]2+ |  | -0.0134333 | -0.017127 |
|  | M638T456_3 | 637.895186 | 7.61 | [729][M+1]2+ |  | -0.0123065 | -0.0171608 |
|  | M640T456_1 | 639.851993 | 7.60 | [737][M]2+ |  | -0.0113178 | -0.0178182 |
|  | M640T456_2 | 639.859472 | 7.61 |  |  | -0.0104075 | -0.0166203 |
|  | M640T456_3 | 639.867468 | 7.61 |  |  | -0.0114184 | -0.0177042 |
|  | M640T457_1 | 639.875145 | 7.61 |  |  | -0.0112872 | -0.0166212 |
|  | M640T457_2 | 639.882824 | 7.61 | [738][M]2+ |  | -0.0113314 | -0.0177269 |
|  | M642T457_1 | 642.324932 | 7.61 | [746][M]2+ |  | -0.0109258 | -0.0179524 |
|  | M642T457_2 | 642.33245 | 7.61 |  |  | -0.0108574 | -0.0172782 |
|  | M642T457_3 | 642.340093 | 7.62 |  |  | -0.0117694 | -0.0173735 |
|  | M642T457_4 | 642.347806 | 7.62 | [747][M]2+ |  | -0.0119698 | -0.0174446 |
|  | M643T457_1 | 642.834867 | 7.61 | [746][M+1]2+ |  | -0.0101979 | -0.0157111 |
|  | M659T465_1 | 659.386497 | 7.75 | [766][M]2+ |  | -0.0117899 | -0.0194691 |
|  | M659T465_2 | 659.394458 | 7.75 |  |  | -0.0120167 | -0.0194397 |
|  | M659T464_1 | 659.402449 | 7.74 |  |  | -0.0128677 | -0.0183468 |
|  | M659T464_2 | 659.410406 | 7.74 | [767][M]2+ |  | -0.012455 | -0.0188563 |
|  | M660T465_2 | 659.90088 | 7.75 |  |  | -0.0130456 | -0.0188385 |
|  | M662T465_6 | 662.379487 | 7.75 | [771][M+1]2+ |  | -0.0119088 | -0.0196229 |
|  | M662T465_7 | 662.387561 | 7.74 |  |  | -0.0121793 | -0.0180809 |
|  | M682T472_1 | 681.907326 | 7.86 | [786][M+1]2+ |  | -0.0127004 | -0.0199883 |
|  | M682T472_2 | 681.915729 | 7.86 | [787][M+1]2+ |  | -0.0125969 | -0.0199104 |
|  | M682T472_3 | 681.924185 | 7.86 | [788][M+1]2+ |  | -0.012637 | -0.0200158 |
|  | M703T479_1 | 703.410169 | 7.98 |  | [M+H+NH3]+ 685.388 | -0.0113493 | -0.0192656 |
|  | M703T479_2 | 703.419118 | 7.98 | [814][M]2+ |  | -0.013806 | -0.0168831 |
|  | M703T479_3 | 703.427714 | 7.98 |  | [M+H+NH3]+ 685.388 | -0.0137803 | -0.0175943 |
|  | M703T479_4 | 703.436658 | 7.98 |  | [M+K+NH3]+ 647.448 [M+Na+NH3]+ 663.421 [M+H+NH3]+ 685.403 | -0.0134555 | -0.0176596 |
|  | M704T479_1 | 703.923621 | 7.98 | [814][M+1]2+ |  | -0.0138542 | -0.0182407 |
|  | M704T479_2 | 703.93245 | 7.98 |  | [M+H+NH3]+ 685.901 | -0.0138344 | -0.0186102 |
|  | M704T479_3 | 703.941564 | 7.98 | [815][M+1]2+ |  | -0.0111573 | -0.0194685 |
|  | M706T479_1 | 705.890313 | 7.98 |  | [M+Na]+ 682.905 [M+H]+ 704.887 | -0.013397 | -0.018718 |
|  | M706T479_2 | 705.899108 | 7.98 |  | [M+Na]+ 682.905 [M+H]+ 704.887 | -0.0134922 | -0.0189003 |
|  | M706T478_1 | 705.907976 | 7.97 |  | [M+Na]+ 682.928 [M+H]+ 704.91 | -0.0135676 | -0.0191987 |
|  | M706T478_2 | 705.916847 | 7.97 |  | [M+Na]+ 682.928 [M+H]+ 704.91 | -0.0136005 | -0.019453 |
|  | M725T485_1 | 725.427537 | 8.09 | [830][M]2+ |  | -0.0132702 | -0.0176122 |
|  | M725T485_2 | 725.436796 | 8.09 |  |  | -0.0136304 | -0.017764 |
|  | M725T485_3 | 725.446062 | 8.09 |  |  | -0.0137952 | -0.0179665 |
|  | M725T485_4 | 725.455263 | 8.08 | [831][M]2+ |  | -0.0130963 | -0.0182164 |
|  | M726T485_1 | 725.937464 | 8.09 | [830][M+1]2+ |  | -0.013518 | -0.014674 |
|  | M726T485_2 | 725.946659 | 8.09 | [831][M+1]2+ |  | -0.0132776 | -0.0156483 |
|  | M728T485_1 | 728.414756 | 8.09 |  |  | -0.0135898 | -0.0171831 |
|  | M728T485_2 | 728.423954 | 8.09 |  |  | -0.0141058 | -0.0177795 |
|  | M750T491_1 | 749.928648 | 8.19 |  |  | -0.0137401 | -0.01693 |
|  | M750T491_2 | 749.938383 | 8.19 |  |  | -0.0140348 | -0.0171043 |
| C18 Negative | M564T584_1 | 564.10466 | 9.74 |  | [M+Cl]- 529.145 [M-H]- 565.121 | -0.0172695 | 0.0153133 |
| C18 Negative | M564T584_2 | 564.111014 | 9.74 |  | [M+Cl]- 529.145 [M-H]- 565.121 | -0.016888 | 0.0158583 |
| C18 Negative | M564T584_3 | 564.117357 | 9.74 |  | [M+Cl]- 529.145 [M-H]- 565.121 | -0.016991 | 0.01577289 |
| C18 Negative | M564T554 | 564.117298 | 9.24 |  |  | -0.0168645 | 0.01486005 |
| C18 Negative | M564T584_4 | 564.123705 | 9.74 |  | [M+Cl]- 529.145 [M-H]- 565.121 | -0.0169758 | 0.0164383 |
| C18 Negative | M566T575_1 | 566.111921 | 9.58 |  | [M+Cl]- 531.146 [M-H]- 567.122 | -0.0183743 | 0.01465833 |
| C18 Negative | M566T584_1 | 566.111993 | 9.74 |  |  | -0.0162502 | 0.01653315 |
| C18 Negative | M566T575_2 | 566.118309 | 9.58 |  | [M+Cl]- 531.146 [M-H]- 567.122 | -0.0184635 | 0.0149058 |
| C18 Negative | M566T584_2 | 566.118378 | 9.74 |  |  | -0.0161048 | 0.01609361 |
| C18 Negative | M606T592_1 | 606.11213 | 9.86 |  | [M+Cl]- 571.146 | -0.0125693 | 0.01342044 |
| C18 Negative | M606T619_1 | 606.112144 | 10.32 |  | [M+Cl]- 571.146 [M-H]- 607.123 | -0.0128963 | 0.01596625 |
| C18 Negative | M606T640_1 | 606.1121 | 10.66 |  | [M+Cl]- 571.15 [M-H]- 607.126 | -0.0127031 | 0.01535215 |
| C18 Negative | M606T592_2 | 606.119207 | 9.86 |  | [M+Cl]- 571.146 | -0.0126471 | 0.01327077 |
| C18 Negative | M606T619_2 | 606.119222 | 10.32 |  | [M+Cl]- 571.146 [M-H]- 607.123 | -0.0130426 | 0.01589101 |
| C18 Negative | M606T640_2 | 606.119177 | 10.66 |  | [M+Cl]- 571.15 [M-H]- 607.126 | -0.0128196 | 0.01541979 |
| C18 Negative | M606T592_3 | 606.126284 | 9.86 |  | [M+Cl]- 571.165 | -0.0127462 | 0.01310416 |
| C18 Negative | M606T619_3 | 606.1263 | 10.32 |  | [M-H]- 607.141 | -0.0131634 | 0.01584598 |
| C18 Negative | M606T640_3 | 606.126248 | 10.66 |  | [M+Cl]- 571.15 [M-H]- 607.126 | -0.0139195 | 0.01688238 |
| C18 Negative | M606T640_4 | 606.133324 | 10.66 |  | [M+Cl]- 571.167 [M-H]- 607.144 | -0.0141463 | 0.01702072 |
| C18 Negative | M606T619_4 | 606.133378 | 10.32 |  | [M-H]- 607.141 | -0.0134093 | 0.01423602 |
| C18 Negative | M606T619_5 | 606.140457 | 10.32 |  | [M-H]- 607.141 | -0.0134335 | 0.01421024 |
| C18 Negative | M606T640_5 | 606.140401 | 10.66 |  | [M+Cl]- 571.167 [M-H]- 607.144 | -0.0143307 | 0.01722117 |
| C18 Negative | M608T592_1 | 608.112714 | 9.86 |  | [M-2H+K]- 571.165 | -0.0131961 | 0.01460066 |
| C18 Negative | M608T627_1 | 608.112681 | 10.46 |  |  | -0.0122196 | 0.01599102 |
| C18 Negative | M608T592_2 | 608.119827 | 9.86 |  | [M-2H+K]- 571.165 | -0.0135163 | 0.01497318 |
| C18 Negative | M608T627_2 | 608.119738 | 10.46 |  |  | -0.0125472 | 0.0184865 |
| C18 Negative | M608T619_2 | 608.119896 | 10.32 |  | [M+Cl]- 573.15 [M-H]- 609.127 | -0.0119532 | 0.01817977 |
| C18 Negative | M608T592_3 | 608.126942 | 9.86 |  | [M+Cl]- 573.161 [M-H]- 609.138 | -0.0136858 | 0.01482927 |
| C18 Negative | M608T627_3 | 608.126853 | 10.46 |  | [M+Cl]- 573.161 [M-H]- 609.138 | -0.0122316 | 0.01653782 |
| C18 Negative | M608T619_3 | 608.12696 | 10.32 |  | [M+Cl]- 573.15 [M-H]- 609.127 | -0.0129627 | 0.01584281 |
| C18 Negative | M608T592_4 | 608.134056 | 9.86 |  | [M+Cl]- 573.161 [M-H]- 609.138 | -0.0139911 | 0.0126923 |
| C18 Negative | M608T627_4 | 608.134018 | 10.46 |  | [M+Cl]- 573.161 [M-H]- 609.138 | -0.0123377 | 0.01602038 |
